# Ontology for Cellular Senescence Mechanisms

**DOI:** 10.1101/2023.03.09.531883

**Authors:** Yuki Yamagata, Tsubasa Fukuyama, Shuichi Onami, Hiroshi Masuya

## Abstract

Although cellular senescence is a key factor in organismal aging, with both positive and negative effects on individuals, its mechanisms remain largely unknown. Thus, integrating knowledge is essential to explain how cellular senescence manifests in tissue damage and age-related diseases. Here, we propose an ontological model that organizes knowledge of cellular senescence in a computer-readable form. We manually annotated and defined cellular senescence processes, molecules, anatomical structures, phenotypes, and other entities based on the Homeostasis Imbalance Process ontology. We described the mechanisms as causal relationships of processes and modeled a homeostatic imbalance between stress and stress response in cellular senescence for a unified framework. HoIP was assessed formally, and the relationships between cellular senescence and disease were inferred for higher-order knowledge processing. We visualized cellular senescence processes to support knowledge utilization. Our study provides a knowledge base to help elucidate mechanisms linking cellular and organismal aging.

---

Multicellular organisms are products of repeated rounds of cell growth and division^1^. After repeated divisions, many cells lose their functionality and stop proliferating. This phenomenon is called cellular senescence and represents a cell fate characterized by stable proliferative arrest in response to various stressors^2^. It is implicated as a fundamental contributor to the aging of individuals and as a contributor to age-related and chronic diseases^2^. Consequently, the use of senotherapy or senolytics to ameliorate aging by eliminating senescent cells has become a research focus^3,4^.

Organismal aging is defined as “a progressive process associated with declines in structure and function, impaired maintenance and repair systems, increased susceptibility to disease and death, and reduced reproductive capacity”^5^. The mechanisms underlying the progressive process of aging are often ascribed to cellular senescence. Both organismal aging and cellular senescence are associated with the disturbance of homeostasis. Aging has been considered the collapse of homeostasis^6^, and cellular senescence is seen as a process that maintains cell viability when a cell can no longer contribute to continued homeostasis via cell division^7^.

Cellular senescence presents both intrinsic advantages and disadvantages for organisms^7^; however, its mechanisms remain largely unknown. Organisms are comprised of complex, heterogeneous, and granular structures with various functions. To understand the aging-related phenotypes of organisms, it is necessary to explain the role of cellular senescence in tissue damage, pathological changes, and aging-related diseases at levels ranging from the cellular to organismal. Clarifying these mechanisms will require expert knowledge in many fields, including molecular biology, cell biology, pathology, pharmacology, clinical medicine, and gerontology. Hence, aging research would benefit from knowledge on the mechanisms of cellular senescence being organized in an interdisciplinary manner.

Various ontologies have been developed for biomedical research. Most notably, the Gene Ontology (GO) has played a major role in providing terms in a machine-readable manner and has contributed to the functional analysis of genes and sharing of knowledge^8^. Developing an ontology that can describe and organize entities related to cellular senescence would significantly promote studies on aging by permitting the integration of relevant data and knowledge.

Here, we aimed to systematize the knowledge of cellular senescence processes in a computer-understandable form by developing an ontology. The ontology was designed to function as a common knowledge base with capacity for higher-order knowledge processing, enabling inference of the relationships between cellular senescence and disease.

## Results

### Ontology development based on HoIP for organizing cellular senescence knowledge

To systematically describe the homeostatic disturbance processes underpinning cellular senescence, we utilized the Homeostasis Imbalance Process ontology (HoIP)^9^ as the foundation for knowledge modeling. HoIP consists of three layers and has some fundamental advantages (Fig. 1a). HoIP refers to the top-level Basic Formal Ontology (BFO)^10^ commonly used in over 100 ontology projects worldwide, defining terms with a domain-independent objective viewpoint at the highest layer. HoIP also defines domain-independent functional processes based on functional ontology^11^, which allows specialized definitions of biological functional processes as subclasses acting to maintain homeostasis. In the intermediate layer, HoIP defines the biological processes essential for maintaining homeostasis. To permit interoperability of the knowledge, terms widely used in biomedical ontology in the OBO Foundry^12^, such as GO, the phenotype ontologies PATO^13^ and HPO^14^, the anatomy ontology UBERON^15^, the cell ontology CL^16^, the Symptom Ontology^17^, the disease ontology DO^18^, the protein ontology PRO^19^, chEBI^20^, and the NCBI taxonomy^21^, were imported into the intermediate layer of the HoIP. The bottom layer relates to the homeostatic imbalance. Here, to deal with the mechanism of homeostasis disturbances, HoIP defines a “homeostatic imbalance course,” which is a process sequence consisting of multiple processes as a subclass of the BFO’s “process.” HoIP is described by Web Ontology Language (OWL)-description logic (DL) language^22^ for both human and machine readability. Thus, we maximized the advantages of HoIP and defined the domain knowledge of cell senescence in the bottom layer. To provide domain-specific knowledge of cellular senescence at the homeostatic imbalance bottom layer, we newly defined terms describing stress responses that act to maintain homeostasis as the functional process in the cellular senescence mechanism (Fig. 1 and Supplementary Table 1). We also defined terms relating to the stresses as functional demands. By challenging the understanding of the causes of processes, we described the mechanism in a unified representation framework as causal relationships and introduced a homeostasis imbalance model. This model illustrates how homeostasis disturbance could lead to tissue damage via cellular senescence, with applications to organismal aging and diseases.

**Fig. 1.**
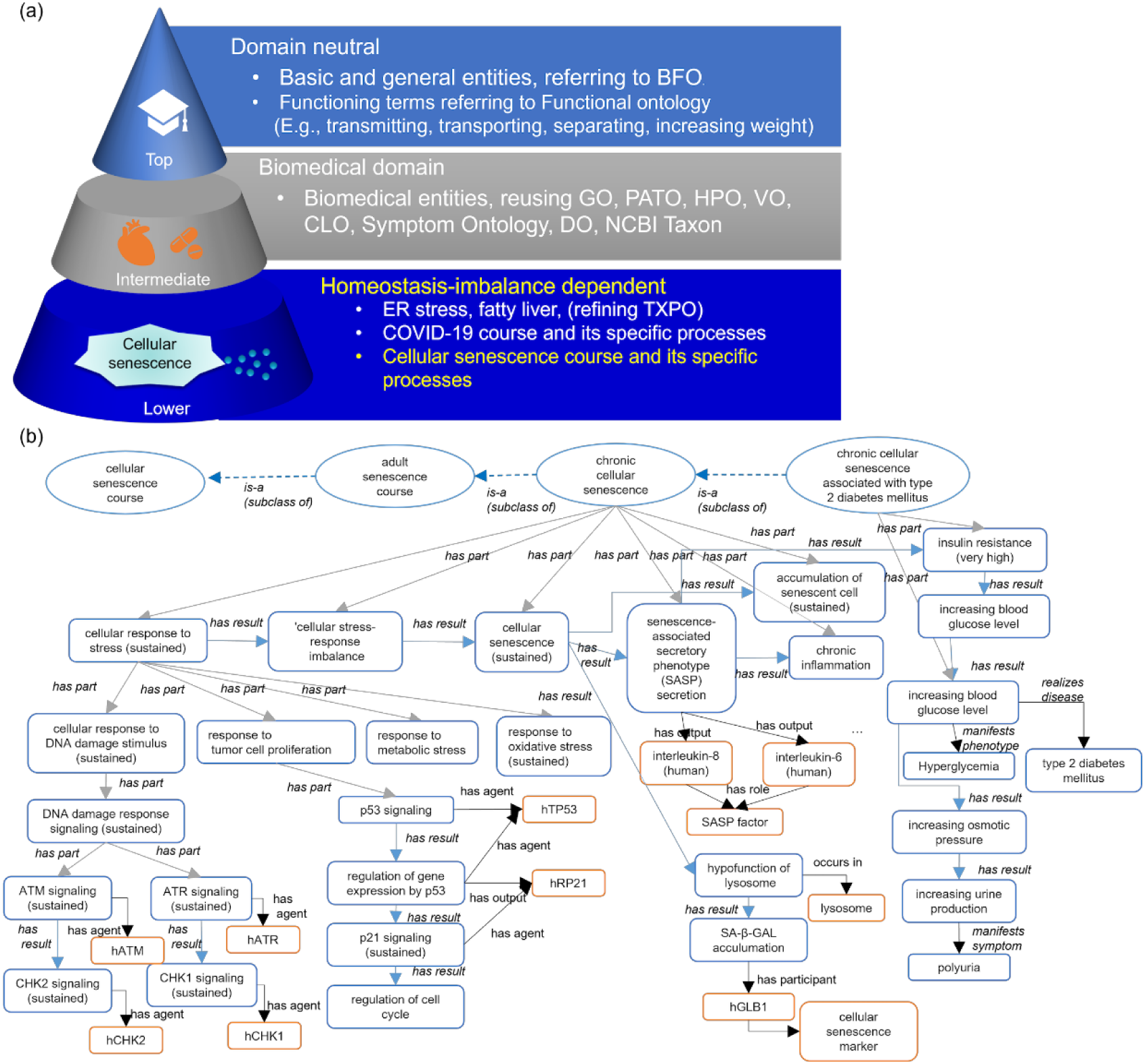
Schematic representation of cellular senescence and HoIP ontology. (a) Overview of the organization of homeostasis imbalance process ontology (HoIP) The HoIP ontology comprises three layers. The top layer is domain-independent and defines basic and general terms that refer to the BFO and functional terms that can be used across domains. The intermediate layer is biomedicine-dependent, and reuses the OBO foundry ontology, such as GO, CLO, HPO, DO, and Symptom Ontology. The lower layer depends on homeostasis imbalance and defines cellular senescence processes and courses. (b) Ontological representation pattern of the cellular senescence course in HoIP. Cellular senescence courses are shown in ellipses. The rounded rectangles are indicated with blue surrounds for processes and occurrents (entities which occur in time) and orange for molecules, roles, and structures. Is-a (subclass of) relationships between courses are indicated by dotted arrows. Blue arrows indicate causal relationships (results), gray for part of the whole (has part), and other relationships are colored black.

### Definition and types of cellular senescence courses to elucidate mechanisms

We first established what processes may occur during cellular senescence. We defined the cellular senescence course, a subclass of “process sequence,” as “the totality of all processes through which cellular senescence is realized.” Subsequently, we classified cellular senescence courses into embryonic and adult cellular senescence courses. In the adult cellular senescence course, since the accumulation of senescent cells can induce pathological processes, we further classified subclasses and defined acute and chronic cellular senescence mechanisms^7^.

**Cellular senescence course**: A course constituting multiple processes that lead to cellular senescence.
  **Embryonic cellular senescence course**: A cellular senescence course that occurs during embryogenesis.
  **Adult cellular senescence course**: A cellular senescence course in adults.
    **Acute cellular senescence course**: An adult cellular senescence course that can cause transient cellular senescence.
    **Chronic cellular senescence course**: An adult cellular senescence course that can result in a sustained cellular senescence process, which may lead to the senescence-associated secretory phenotype (SASP), and chronic inflammation due to senescent cell accumulation.

In addition, we defined chronic cellular senescence as being associated with type 2 diabetes mellitus because it is related to chronic diseases in the elderly (Fig. 1b).

### Computational representation of cellular senescence knowledge

#### 1. Annotated Information for human experts

HoIP is based on OWL, which enables reasoning by description logic. However, it is difficult for domain experts to understand the content. Therefore, in line with HoIP development policy, information for human domain experts was provided using OWL annotation properties. All definitions of entities involved in cellular senescence were newly described in natural language. In addition, entities were manually annotated from textbooks, reviews, and journal articles by human experts. Moreover, to enable researchers to follow the evidence trail behind annotation, the description related to the annotation was extracted from abstracts, original texts, and figure legends. Moreover, reference information (PubMed ID (PMID)) was assigned (Fig. 2 (a)) as an external cross-reference. We focused on human cellular senescence, so molecular entities involved in cellular senescence processes were annotated manually, referred to as the Protein Ontology, and we added cross-references to the HUGO Human Gene Nomenclature Committee (HGNC)^23^. Mouse orthologous genes were also annotated, if available, from Protein Ontology. It should be noted here that data using the annotation properties are supplementary information for the machine and do not affect the inference. In other words, our descriptions distinguished between cellular senescence knowledge in a human-readable form for human experts from that used for computer processing for ontological reasoning.

#### 2. Description and formalization for computer processing

##### (1) Cellular aging-specific processes and related entities

OWL-DL is designed to enable logical computational processing using object properties. Therefore, we described the characteristics of each entity and its relationships with other entities using object properties. Parent classes were also identified. Fig. 2 (b) shows a representation of the entity *SASP secretion* (HOIP:0060102) newly defined in the HoIP using OWL object properties by Protégé software as an example of a process specific to cellular senescence. A process can have sub-processes; for example, SASP secretion has *IL-6 signaling* (HOIP:0060639) as a sub-process using the object property ‘has part.’ Furthermore, if a process can cause some other process, we described the causal relationship (using the ‘has result’ relation). Fig. 2(b) shows *chronic inflammation* (HOIP:0060113) due to sustained SASP secretion. In the case of SASP secretion, besides chronic inflammation, it can lead to multiple results, such as *fibrosis* (HOIP:0060112) and *negative regulation of tissue regeneration* (HOIP:0060182).

This process can be expressed by the following equation (some parts are omitted): senescence-associated secretory phenotype (SASP) secretion in chronic cellular senescence

⊆(senescence-associated secretory phenotype (SASP) secretion in adult cellular senescence

⋂ ∃has part. CXCL8 signaling

⋂ ∃has part. IL-6 signaling

⋂ ∃has output. interleukin-6 (human) ⋂ (∃has role. SASP factor)

⋂ ∃has output. interleukin-8 (human) ⋂ (∃has role. SASP factor)

⋂ ∃has result. chronic inflammation

⋂ ∃has result. fibrosis

⋂ ∃has result. hypofunction of keeping stem cell niche

⋂ ∃has result. negative regulation of tissue regeneration

⋂ ∃has result. positive regulation of carcinogenesis

⋂ ∃has result. positive regulation of cellular senescence in neighboring cell

⋂ ∃occurs in. senescent cell

⋂ ∀has context. chronic cellular senescence)

We used the ‘occurs in’ relationship to describe the location of the process. For example, SASP secretion occurs in senescent cells. In the SASP secretory process, *interleukin 6* and *interleukin-8* are secreted and were represented by the use of the ‘has output’ relation. We identified the role for each molecule in processes as context, such as the ‘SASP factor’ role, using the ‘has role’ property. Furthermore, since symptoms are essential in clinical medicine, the ‘manifests symptom’ relationship was used. For example, the process ‘*increasing urine production* (HOIP:0060496)’ manifests as ‘*polyuria* (SYMP_0000565).’ In a case where a process manifested a disease, the ‘realizes disease’ relationship was used to describe diseases, such as *type 2 diabetes mellitus* (DOID:9352) (Fig. 1 (b)). With these properties, we could infer related symptoms and diseases by DL-query.

##### (2) The cellular senescence course

Supplementary Fig. S1 shows an example of a representation of the *chronic cellular senescence course* (HOIP: 0060195) using Protégé. The processes defined above (1) were enumerated using the ‘part of’/‘has part’ relation as the object property for each course of cellular senescence, and they were represented as belonging to the course, that is, as constituting a course. The subclasses of the course inherited the processes of the course of the superclass, specialized them, or added new processes. For example, *cellular senescence* (HOIP:0060129) in the cellular senescence course was specialized to *cellular senescence (sustained)* (HOIP:0060240) in the chronic senescence course, and processes such as *accumulation of senescent cell (sustained)* (HOIP:0060308) and *chronic inflammation* (HOIP:0060113) were added to the components. This equation is presented in (Supplementary Information 1).

#### 3. Unified representation model of homeostatic imbalance in cellular senescence: causal networks in cellular senescence mechanisms

Because each process in the cellular senescence course has a causal relationship, a network (causal network) consisting of linked causal relationships among multiple processes was generated by linking them computationally. Here, we applied the imbalance balance model to the network and provided a unified description by representing the imbalance between functional demand and the functioning process^24,25^. As a result of the formalization described above, a unified descriptive framework model was constructed to describe the progression of the cellular senescence mechanism as follows: 1) stress (functional demand process), 2) stress response as the functioning process to maintain homeostasis, 3) homeostatic imbalance, and 4) outcome, which is cellular senescence, as summarized in Fig. 3 (a).

**Fig. 2.**
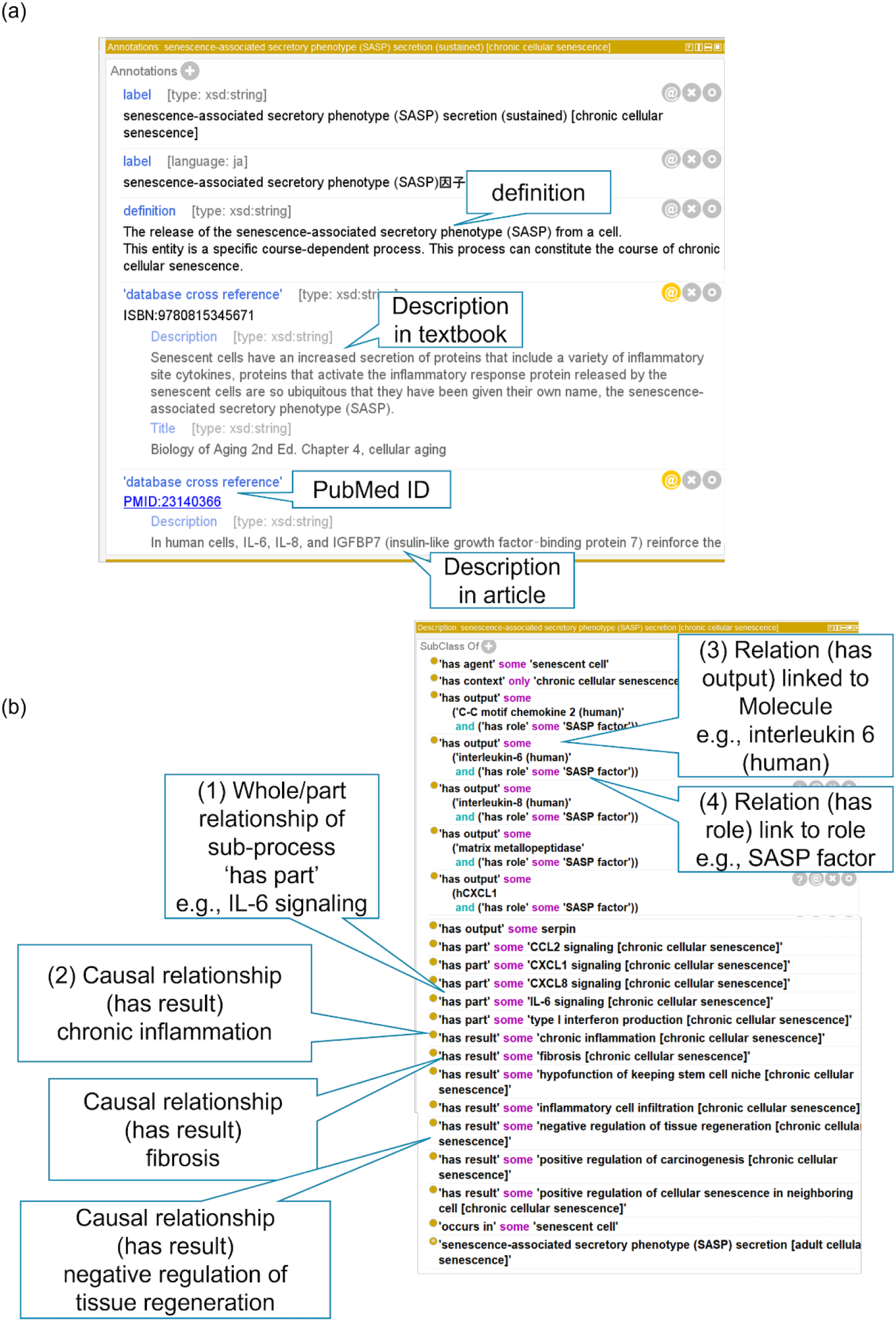
Example of computational representation of senescence-associated secretory phenotype (SASP) secretion (HOIP:0060102) with the Protégé software. (a) A screenshot from Protégé for an example of a description using the annotation properties. Descriptions by natural language are given for the “definition”, “Description” from textbooks or articles, and links to external references with “PubMed ID” are shown by balloons. (b) Example of a description using object properties of the process. The major entities involved in the process are indicated by balloons. In this example, relations to process as Senescence-associated secretory phenotype (SASP) secretion, (1) whole/part (*‘*has part’) relation linked to sub-process as IL6 signaling, (2) causal relation (‘has result) linked to chronic inflammation, (3) relation (has output) linked to a molecule such as interleukin-6 (human) (4) as a “role”, SASP factor.

**Fig. 3.**
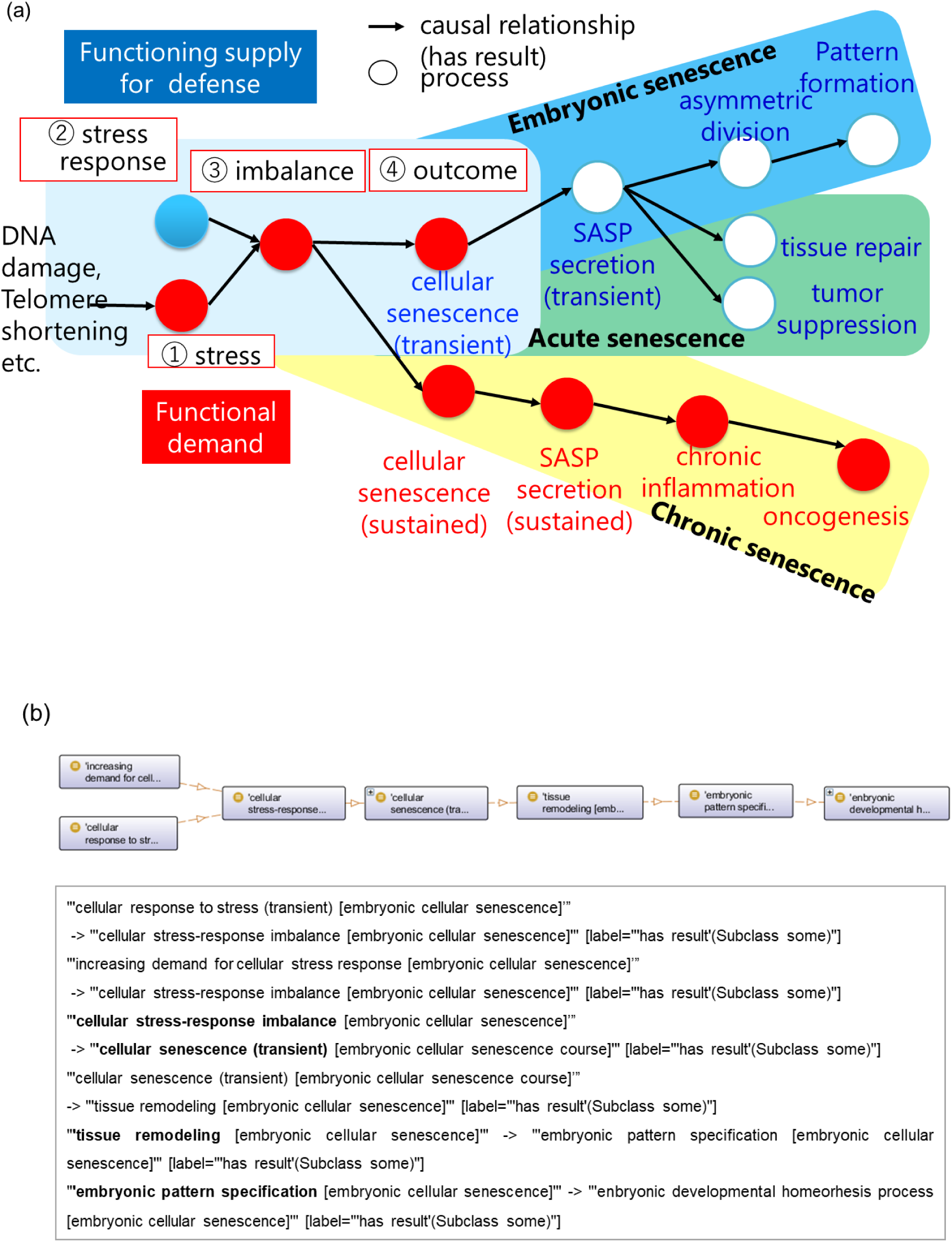
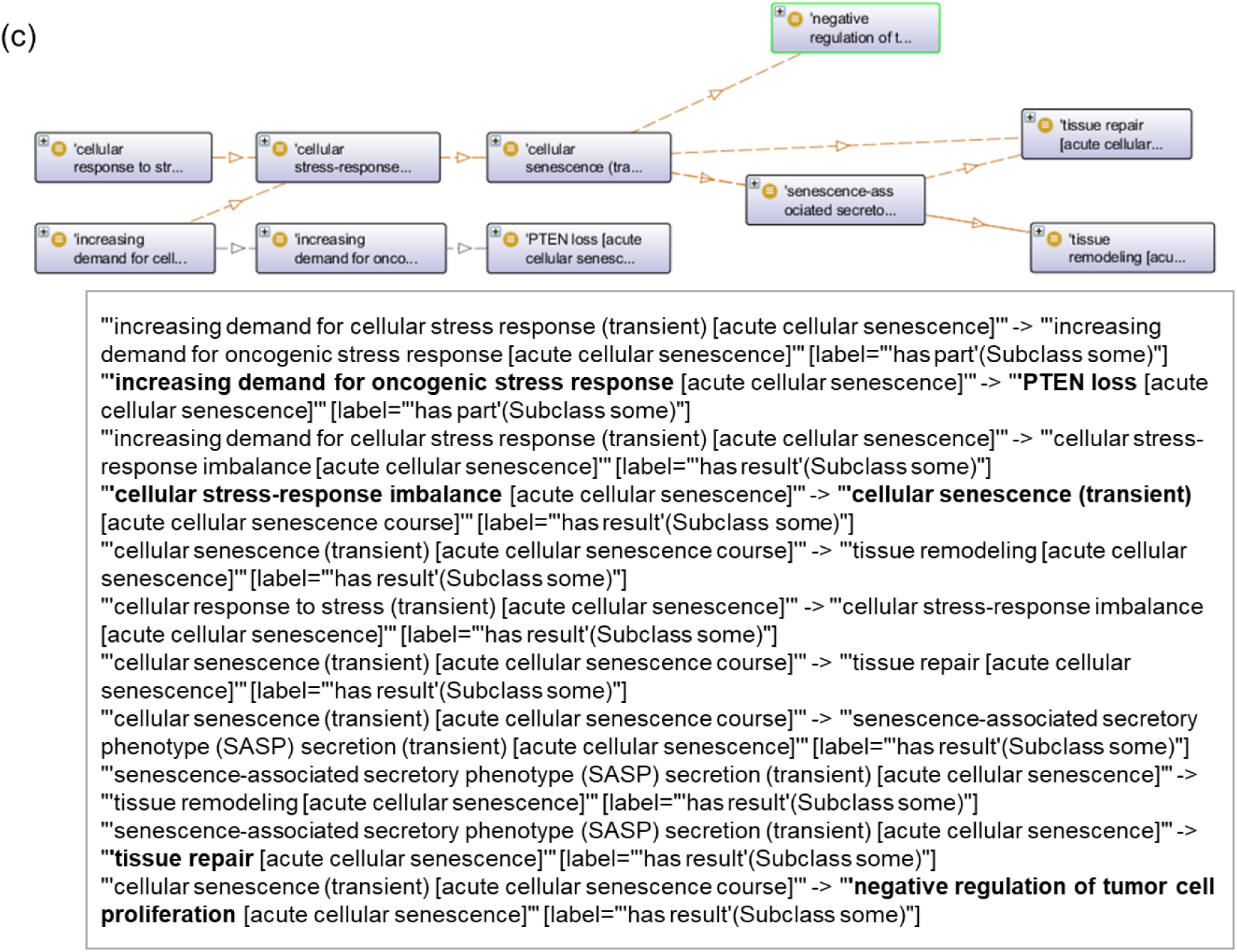
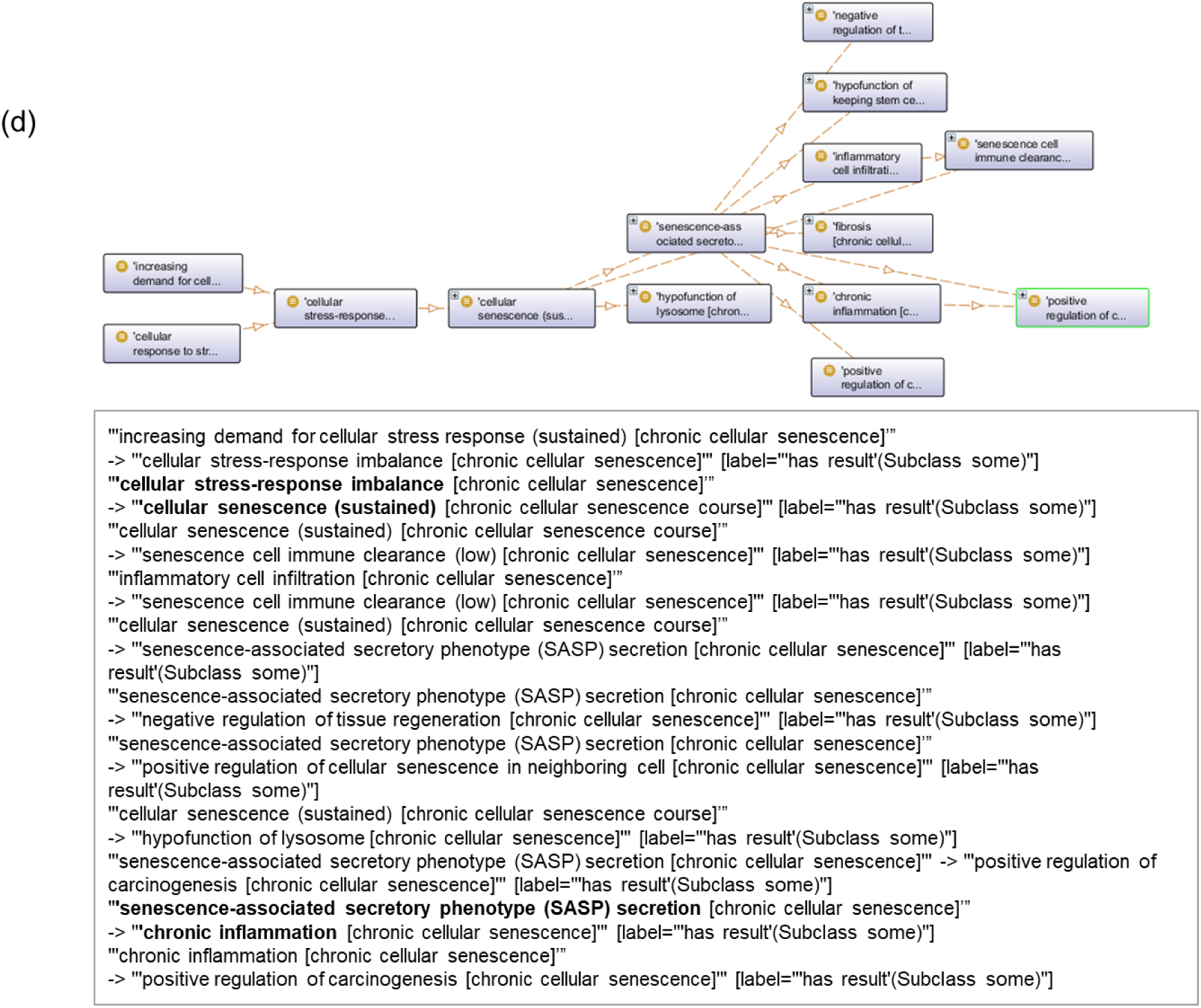
A unified representation of cellular senescence courses. (a) An overview of the unified representation of cellular senescence courses. The blue terms indicate processes that can be caused by transient cellular senescence, and the red terms indicate processes that can be caused by sustained cellular senescence. The squares indicate the basic components of homeostatic imbalance: (1) stress, (2) stress response, (3) imbalance, and (4) outcome. The embryonic cellular senescence course is shown in a blue round box, the acute cellular senescence course is shown in a green round box, and the chronic cellular senescence course is shown in a yellow round box. (b) The upper figure shows a screenshot of OntoGraf in Protégé displaying the imbalance model in the embryonic cell aging course, with ‘Arc Types’ selected to be causal relationships (“has result”) for the cell aging process. Lower indicates a partial excerpt of the causal relationships (shown in “->”) using the ‘export DOT’ function from OntoGraf. In embryonic cellular senescence, the cellular-stress imbalance could result in cellular senescence (transient). Consequently, tissue remodeling could cause pattern formation specification. (c) The upper figure shows causal relationships introduced by the imbalance model in acute cellular senescence course using OntoGraf in Protégé. In the case of acute cellular senescence, there is increasing demand for the oncogenic stress response, which has the subprocess PTEN loss, and the imbalance could lead to cellular senescence (transient), tissue repair, and negative regulation of tumor proliferation. (d) The upper figure shows causal relationships introduced by the imbalance model in chronic cellular senescence course using OntoGraf in Protégé. In chronic cellular senescence, imbalance could cause persistent cellular senescence, which might lead to SASP secretion, which could cause chronic inflammation.

Using the OntoGraph API of Protégé, our model demonstrated that in the embryonic cellular senescence course and the acute cellular senescence course, the homeostatic imbalance could lead to transient cellular senescence, as shown in Fig. 3 (b), (c). In embryonic cellular senescence, our model showed an imbalance in stress as a demand for asymmetric division, and the response could result in transient cellular senescence. Consequently, tissue remodeling can result in embryonic pattern specification (Fig. 3 (b)). In the case of acute cellular senescence, our model described that as part of the increasing demand for a cellular stress response, there might be an increasing demand for the oncogenic stress response, which has a further subprocess, PTEN loss. Furthermore, the imbalance between stress and its response as a defense function could lead to transient cellular senescence, which might lead to negative regulation of tumor proliferation (Fig. 3 (c)). In contrast, in chronic cellular senescence, the imbalance model showed that persistent cellular senescence, leading to SASP secretion, could cause further chronic inflammation (Fig. 3 (d)).

In summary, our ontology and the imbalance model enable a unified representation of the causal networks of cellular senescence.

### Assessment of Ontology Quality

To assess the quality of the constructed ontology, a mechanical approach^26^ was first taken. We used the ontology reasoner HermiT^27^ and ELK^28^ in Protégé for formal verification. The results showed no violations of the domain or range of the property class hierarchy, and there was consistency in terms of formality. Next, to validate the content, we inferred whether DL queries could adequately obtain the causal relationships between processes related to each cellular senescence course as use cases.

#### (1) Validation of the causal network of p21 signaling

The mechanism of cellular senescence involves p16 and p21 signaling^3,7,29^. Therefore, we validated the causes of p21 signaling using ontology reasoning as a case study. We used a transitive property, owl:TransitiveProperty, for the causal relation (‘has result’), which infers not only direct causal relations but also indirect causal relations. The results showed that in *adult cellular senescence* (HOIP:0060315), for example, *p21 signaling* (HOIP:0060325) could occur via *ATM* signaling (HOIP:0060110), *p53 signaling* (HOIP:0060055), and regulation of gene expression by p53 (Supplementary Fig. S2 (a)). On the other hand, in the *embryonic senescence course* (HOIP:0060267), the results indicated that p21 signaling could occur via *TGF beta signaling* (HOIP:0060269), *SMAD signaling* (HOIP:0060290) and *regulation of gene expression by SMAD* (Supplementary Fig. S2 (b)). Thus, our model illustrates that the differences between courses can be represented by different paths in the causal networks.

#### (2) Validation of causal network including part of the whole relation

Next, we examined whether the causes were adequately described, including the systematic relationship between the whole body and its parts. We validated the causes of p21 signaling in the adult cellular senescence course, including part-whole processes.

We used the OWL2 property chain axiom and defined property “has part result” as the sub-property of the “has result” property: ‘has part’ ∘ ‘has result’ ⊆ ‘has result’

This property allows causal relationships, including whole-part relationships, which are not directly described. The results showed that *ATR signaling* (HOIP:0060140) and *ATM* signaling (HOIP:0060110) upstream of *p21 signaling* (HOIP:0060325) are sub-processes of the *DNA damage response* signaling (HOIP:0060337) (Fig. 4 (a), (b), (c)). Thus, the present study illustrates that our description can explain how the processes involved in cellular senescence, including the whole-part relationships, can be affected.

**Fig. 4.**
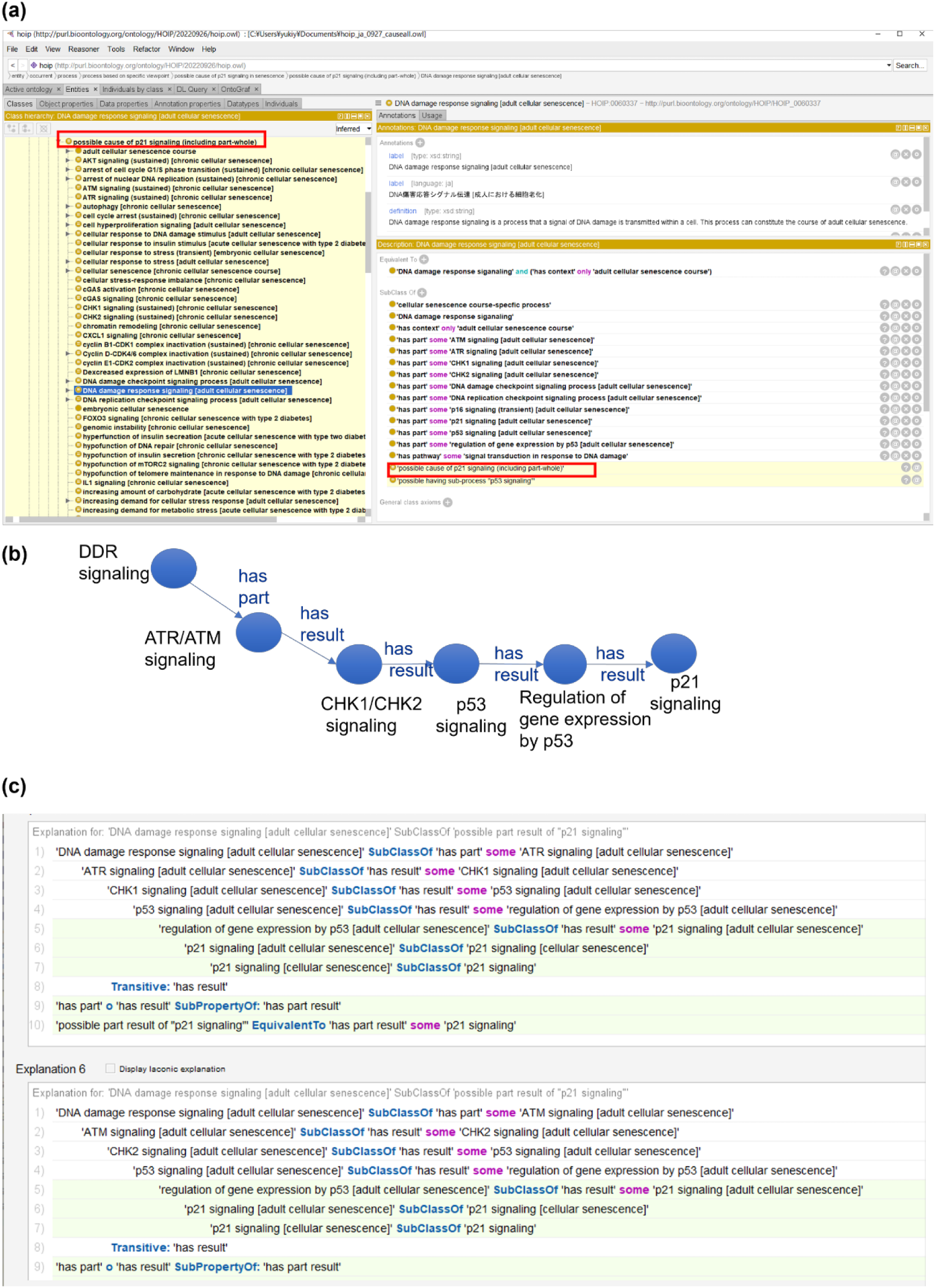
Validation of causal networks including part of whole (‘has part’) relations of processes by ontology reasoning tool ELK in Protégé. (a) A screenshot for a list of the results of causal inference of the ontology reasoner ELK in Protégé software, including part of whole relations of processes and p21 signaling (HOIP:0060325) in the adult cellular senescence course used in the ELK reasoner in Protégé. The upper red frame shows the possible cause of p21 signaling [adult cellular senescence] for validating causal networks. Inferred information is shown on a yellow background (lower red frame). The DNA damaging response signaling (HOIP:0060337) is inferred to be a subclass of possible cause of p21 signaling [adult cellular senescence]. (b) Schematic representation of the results of the inference. Each node shows the process, and each edge shows the relationship (has part/has result) between processes. (c) Validation of causal relation network of DNA damage response signaling (HOIP:0060337) by the ontology reasoning tool ELK in Protégé. The results showed DNA damage response signaling (HOIP:0060337) has subprocesses, ATR signaling (HOIP:0060140) (upper figure) or ATM signaling (HOIP:0060110) (lower figure), which could cause CHK1 signaling (HOIP:0060144) (upper figure)/CHK2 signaling (HOIP:0060145) (lower figure). This could then result in p53 signaling (HOIP:0060055), regulation of gene expression by p53 (HOIP:0060065), and p21 signaling (HOIP:0060325).

We also confirmed that SPARQL queries (Supplementary Information 2) could acquire the processes of chronic cellular senescence (HOIP:0060195), including telomere shortening, *p16 signaling* (HOIP:0060294), *p53 signaling* (HOIP:0060321), and *senescence-associated heterochromatin focus (SAHF) formation* (HOIP:0060162).

### Causal inference of disease-related processes

As a case study, we used our ontology to infer the possible causal relationships between the cellular aging process and type 2 diabetes mellitus based on a description logic (DL) query.

#### (1) Exploring the relationship between cellular senescence and type 2 diabetes mellitus

The relationship between cellular senescence and disease remains a compelling question. Ontology-based inference is expected to support the discovery of causal networks between cellular senescence and disease through computational processing. In this study, we used ELK to explore the relationship between cellular senescence and type 2 diabetes mellitus, a chronic disease of aging. After manually annotating known mechanisms associated with type 2 diabetes mellitus, we inferred paths computationally using OWL transitivity to find latent paths via indirect causal relations. Fig. 5 (a) shows a list of processes involved in the causes of *insulin resistance* (HOIP:0060427). One hypothesis was that *telomere shortening* (HOIP:0060431) could cause *Type B pancreatic cell exhaustion* (HOIP:0060453), resulting in *positive regulation of SASP secretion* (HOIP:0060455) and *SASP secretion* (HOIP:0060432), which might lead to *insulin resistance* (HOIP:0060427). Other processes involved included *FOXO signaling* (HOIP:0060504) and *TLR4 signaling* (HOIP:0060512) (Fig. 5 (b)).

#### (2) SASP factors and cross-mechanism causal search: relationship to COVID-19 severity mechanisms

During chronic cellular senescence, the accumulation of senescent cells can cause SASP secretion and lead to *IL-6 signaling* (HOIP:0060639) and CXCL8/*IL-8 signaling* (HOIP:0060640). Therefore, we inferred the possible effects of IL-6 and IL-8 signaling across courses in HoIP, finding that inflammation-related processes are common to the cellular senescence course and to COVID-19. Moreover, the inference also showed that the possible effects of IL-8 and IL-6 signaling included processes associated with severe COVID-19, such as *microvascular dysfunction* (HOIP:0040768) and *thrombus formation* in lung (HOIP:0038042) (Fig. 5(c)).

**Fig. 5.**
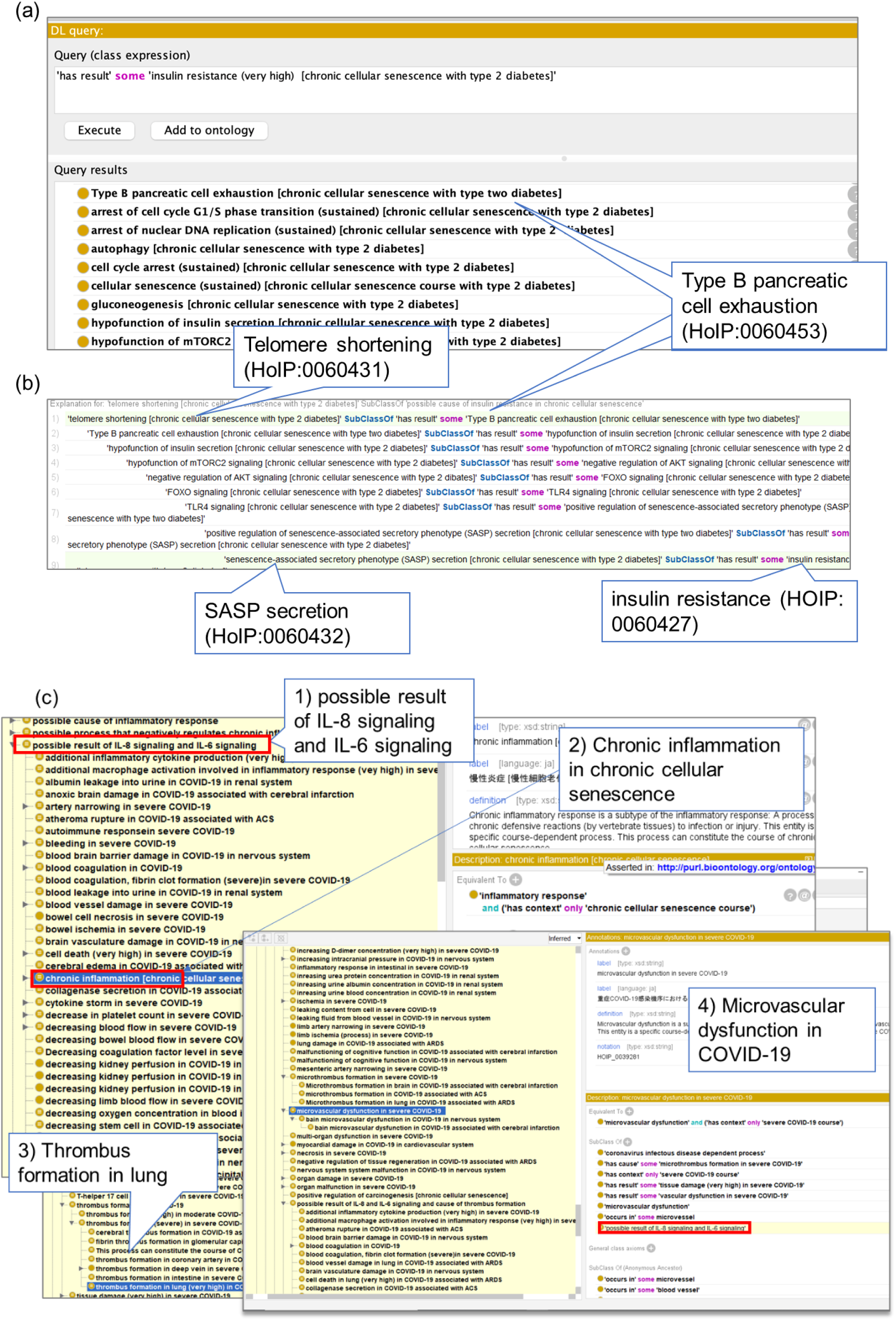
Example of causal inference relevant to diseases in the Protégé software. (a) Inference results in DL query. Upper panel shows the query for the search of causal relationships (has result) of insulin resistance (HOIP:0060427) in chronic cellular senescence associated with type 2 diabetes mellitus (HOIP:0060423). The lower results show the possible cause of the processes by computational reasoning, for example, Type B pancreatic cell exhaustion (HOIP:0060453 (shown in the balloon)). (b) Inference shows that telomere shortening (HOIP:0060431), indicated in the balloon, might cause insulin resistance (HOIP:0060427) via positive regulation of senescence-associated secretory phenotype (SASP) (HOIP:0060455) and senescence-associated secretory phenotype (SASP) secretion (HOIP:0060432) (balloon). (c) Results of causal inference of IL-6 signaling (HOIP:0004327) and CXCL8 (IL-8) signaling (HOIP:0041903) across courses used in the ELK reasoner in Protégé. The upper red frame shows 1) the possible result of IL-6 signaling (HOIP:004327) and CXCL8 signaling (HOIP:0041903) across courses for the validation of causal networks. Inferred information is shown in a yellow background, and 2) chronic inflammation in chronic cellular senescence (HOIP:0060113) and 3) thrombus formation in the lung (very high) in COVID-19 associated with ARDS (HOIP:0041811), and 4) microvascular dysfunction in severe COVID-19 (HOIP:0039281) is inferred as the possible result of IL-8 and IL-6 signaling. The red frame indicates that the microvascular dysfunction term is classified as a possible result of IL-8 and IL-6 signaling by inference.

### Visualization of the cellular senescence causal network

The visualization of ontologies is expected to have various applications. Ontologies are often published as knowledge graphs in RDF format, where a SPARQL query retrieves the necessary data against the SPARQL endpoint, which is a database that stores RDF data. However, it is difficult for senescence specialists to handle complex SPARQL queries. Therefore, in this study, we visualized cellular aging mechanisms using Cytoscape^30^ so that specialists in aging who are unfamiliar with ontologies and SPARQL queries could easily use cellular senescence knowledge in ontologies. In this study, we used SPARQL queries as the background process. The relationships were visualized in an easy-to-understand manner by converting the query results into nodes and edges of a graph using the CyREST API (Fig. 6, see Methods and Supplementary Fig. S3).

**Fig. 6.**
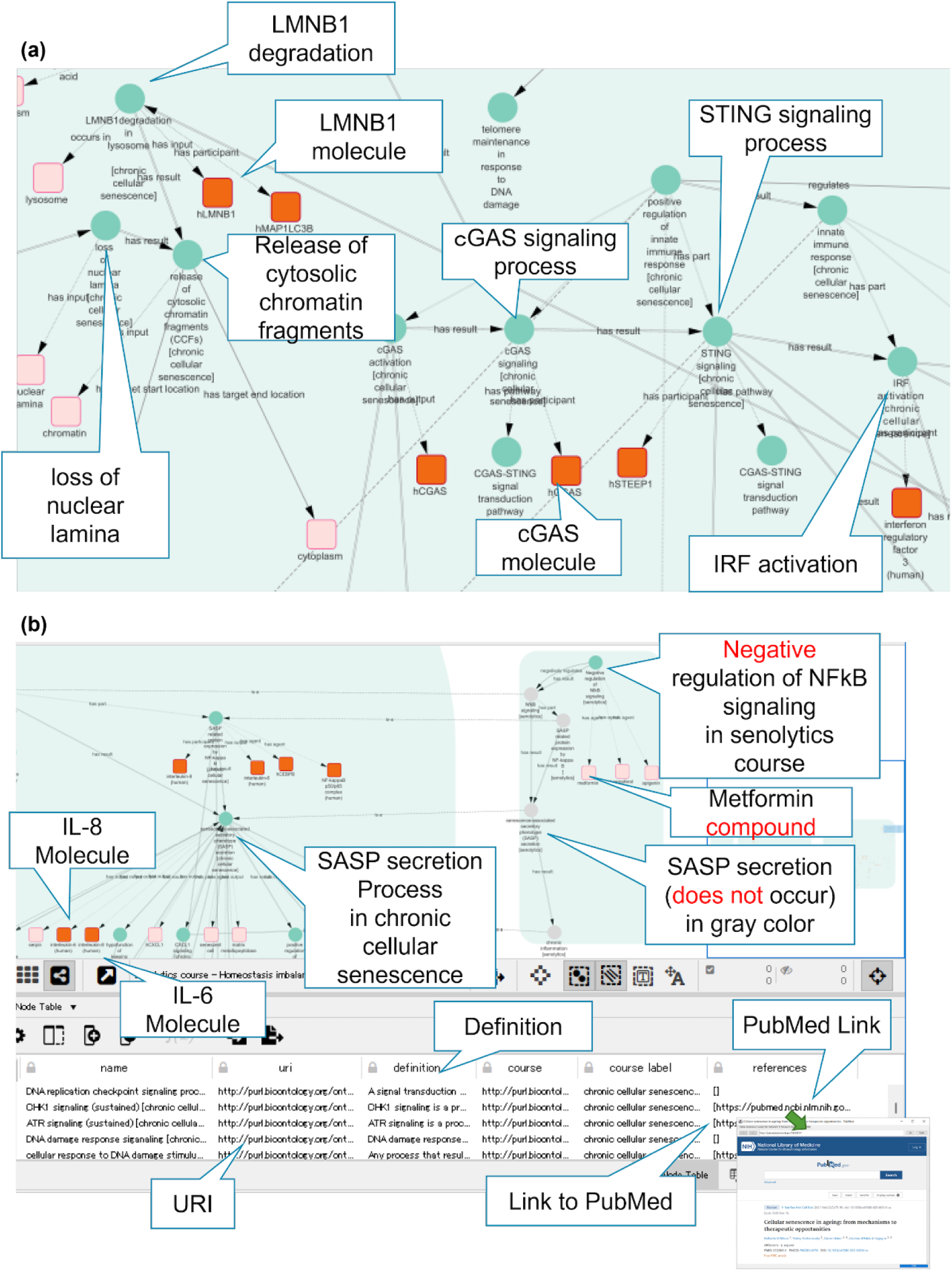
Cytoscape visualization of the cellular senescence course. The graph visualizes causal networks between processes and molecules involved in cellular senescence. Nodes denoted with circles indicate processes, and squares indicate material entities. The red color indicates human molecules. Balloons show explanations of visualized components. (a) Part of a screenshot visualizing the chronic cellular senescence course from LMNB1 degradation to IRF activation using Cytoscape. In the process *LMNB1 degradation in lysosome* (HOIP:0060132), the molecule *LMNB1* (PR:P20700) (red color) is visualized. The downstream visualization showed that the degradation of LMNB1 could cause the *release of cytosolic chromatin fragments (CCFs)* (HOIP:0060190) (balloon), leading to *cGAS signaling* (HOIP:0060172) and *STING signaling* (HOIP:0060179), which might result in *IRF activation* (HOIP:0060183). (b) Part of the screenshot to visualize the process inherited from the super-class which was represented by the “is-a” edge (dot line) across super-class and sub-class courses. In the senolytics course (HOIP:0060418), NFKB signaling in chronic cellular senescence is inherited, and negative regulation of NFKB signaling (HOIP:0060417) by drugs such as metformin (CHEBI_6801) in the senolytics course could cause no NFKB signaling. As a result, downstream, no SASP secretion might occur (right side, gray node). In the lower table, information from the ontology is shown, such as the definition, and a PubMed link.

Our visualization model allows users to search for molecules and processes. We confirmed that Cytoscape could create subgraphs of cellular senescence by simply selecting the types of edges by filtering and could also extract and visualize the neighboring nodes of processes and molecules of interest. In addition, tables can access ontology information in ontology repository sites such as BioPortal^31^ from the URI information of each node. In HoIP, many processes were defined as a sub-class of GO, and the GO URI enabled access to the external GO site from Cytoscape. For each process, via external links to PubMed, a literature search of PubMed was displayed. An example of a part of the visualization of the chronic cellular senescence course is shown in Fig. 6. In the process *LMNB1 degradation in lysosome* (HOIP:0060132), *LMNB1* (PR_P20700) could be visualized. Downstream analysis showed that degradation could cause the *release of cytosolic chromatin fragments (CCFs)* (HOIP:0060190), leading to *IRF activation* (HOIP:0060183) via cGAS-STING signaling (*cGAS signaling* (HOIP:0060172), *STING signaling* (HOIP:0060179)) as an innate immune response (Fig. 6 (a)).

For visualization, the process inherited from the super-class was represented by the “is-a” edge across the super-class and sub-class courses. For example, NFKB signaling in chronic cellular senescence is inherited, and *negative regulation of NFkB signaling* (HOIP:0060417) by drugs such as *metformin* (CHEBI_6801) in the *senolytics course* (HOIP:0060418) could inhibit NFKB signaling. Consequently, no SASP secretion might occur downstream (Fig. 6 (b), right side, grey node).

In this way, we demonstrated that information on cellular senescence was presented graphically, with the nodes corresponding to processes and molecules. Other information about cellular senescence in HoIP is presented in tables, allowing the user to navigate literature and ontologies.

Because users would otherwise need to install Cytoscape, we made the visualization available on NDEx for easy access on the web (https://www.ndexbio.org).

## Discussion

In this study, we developed an ontology to establish a knowledge base focused on cellular senescence. We systematized a wide variety of knowledge relating to cellular senescence dispersed in textbooks and articles from multiple domains into a consistent form that can be processed by computers. By applying HoIP as the fundamental tool for knowledge management, we expect to contribute to both theoretical and practical aspects such as senolytics and senotherapy in drug development. Since major biomedical ontologies such as GO, UBERON HPO, and ChEBI have been referred to in HoIP, the ontology will accelerate interoperability and further accumulation of knowledge.

Knowledge representation based on a unified description by the homeostasis imbalance model enabled us to explain some differences and commonalities among cellular mechanisms. Our model showed that the common outcome of embryonic and acute cellular mechanisms is the transient occurrence of cellular senescence, which results in life-supporting activities, as shown in Fig. 3 (b, c). In embryos, homeostatic imbalance could lead to transient cellular senescence, possibly leading to tissue remodeling and pattern formation during embryonic development (Fig. 3 (b)). Although homeostasis is a static state, disturbance of homeostasis might induce so-called homeorhesis, as proposed by Waddington^32^, which leads to a dynamic equilibrium state. In the case of acute cellular senescence, the imbalance between the stress of activation of oncogenes and defensive responses could lead to transient cellular senescence as well as embryonic cellular senescence, resulting in negative regulation of tumor proliferation (Fig. 3 (c)). Thus, our ontology demonstrated that embryonic and acute cellular senescence might maintain life by bringing about stability at the tissue level, while providing dynamism at the cell level through temporary, local cellular senescence.

In contrast, in chronic cellular senescence, chronic inflammation can result, leading to tissue damage, and potentially, carcinogenesis. Our results indicate that chronic cellular senescence might lead to the collapse of the system in an organism and ultimately death.

Thus, our model reflects biological phenomena in a computational processable way, and indicates that different types of cellular senescence can be either advantageous or disadvantageous to life.

One of the difficulties in sharing knowledge is that each expert has a different knowledge level depending on his/her background. Ontology can solve this problem by providing both general and domain-specific knowledge. The hierarchical ontology structure helps identify commonalities and differences among the diversity of expert knowledge. Furthermore, the ontology can be verified using description logic. In this study, the description of HoIP is based on OWL-DL, and we can formally verify and guarantee consistency using ontology reasoning tools. Description logic is also powerful for causal reasoning. In this study, we use the OWL transitivity axiom. We found multiple paths leading to insulin resistance in chronic cellular senescence associated with type 2 diabetes mellitus. The insulin pathways have also been reported to be relevant to longevity^33^. We are planning to further investigate the effects of insulin on metabolism, the endocrine system, starvation, energy production, and growth, and how these mechanisms are conserved across species.

The importance of aging research has been accelerating, as the risk of severe illness with COVID-19 increases with age. Ontological reasoning will contribute to the elucidation of the crosstalk between COVID-19 and chronic cellular senescence. In this study, we inferred that IL-6 and IL-8 are inflammatory substances in COVID-19 and as SASP factors in cellular senescence (Fig. 5c). Thus, molecules with multiple roles in the body may be a factor in disease severity in the elderly. In support of this, a recent review reported that SARS-CoV-2 induces hyperinflammation in SASP^34^. It was proposed that a propensity for senescence-governed immune escalation, which can be related to aging or chronic disease, can also be acutely triggered by SARS-CoV-2 infection and lead to severe disease. Hence, the results of this study inform the commonalities between COVID-19 and cellular senescence. Here, for the processes that constitute the cellular senescence course and the COVID-19 infectious course, the common processes in both can be computationally obtained by referring to the upper classes of terms as generalized processes using the hierarchical tree of the HOIP ontology. As shown in Table 1, many common processes are related to innate immunity and inflammatory responses.

**Table 1.** Common processes in chronic *cellular senescence course* (HOIP:0060195) and COVID-19 infectious courses: *COVID-19 infectious course* (HOIP:0037002) or *severe COVID-19 infectious course* (HOIP:0039222) or *COVID-19 course associated with acute respiratory distress syndrome* (HOIP:0036006)

| Process URI | Process |
| --- | --- |
| HOIP:0000102 | increasing demand for response to oxidative stress |
| HOIP:0000192 | inflammatory response |
| HOIP:0000219 | fibrosis |
| HOIP:0000540 | inflammatory cell infiltration |
| HOIP:0000761 | NFKB signaling |
| HOIP:0002073 | organ malfunction |
| HOIP:0002361 | mitochondrial dysfunction |
| HOIP:0004327 | IL-6 signaling |
| HOIP:0004451 | CCL2 signaling |
| HOIP:0004657 | IL1 signaling |
| HOIP:0036193 | IRF3 activation |
| HOIP:0040446 | decreasing stem cell |
| HOIP:0040448 | negative regulation of tissue regeneration |
| HOIP:0041903 | CXCL8 signaling |
| GO:0002532 | production of molecular mediator involved in inflammatory response |
| GO:0032606 | type I interferon production |
| GO:0043065 | positive regulation of apoptotic process |
| GO:0045089 | positive regulation of innate immune response |
| GO:0050729 | positive regulation of inflammatory response |
| GO:1903409 | reactive oxygen species biosynthetic process |
| NCIT:C16507 | DNA damage |

HOIP: http://purl.bioontology.org/ontology/HOIP
E.g., HOIP:0000102 http://purl.bioontology.org/ontology/HOIP/HOIP_0000102
GO: http://purl.obolibrary.org/obo/GO.owl
E.g., GO:002532 http://purl.obolibrary.org/obo/GO_0002532
NCIT: http://purl.obolibrary.org/obo/NCIT.owl
E.g., NCIT: C\_16507 http://purl.obolibrary.org/obo/NCIT_C_16507

Furthermore, a generic causal network can be generated across multiple courses using generalized processes. For example, in the chronic cellular senescence course, SASP secretion and its subprocess, IL-8 signaling, might lead to chronic inflammation. On the other hand, in the COVID-19 course, thrombus formation can occur through multiple pathways, from IL-8 signaling to neutrophil extracellular trap formation and NETosis. A generalized causal network of IL-8 signaling as the subprocess of SASP secretion *→* neutrophil extracellular trap formation/ NETosis *→* thrombus formation can be derived. Furthermore, IL-8 signaling can result from NFKB signaling via cGAS-STING signaling in chronic cellular senescence due to increasing extrachromosomal telomere repeat DNA from telomere shortening or from the release of cytoplasmic chromatin fragments due to the loss of nuclear lamina. Therefore, even if the virus is no longer in the body, thrombus formation may still occur if cellular senescence-derived IL-8 signaling occurs (Table 2). A review reported that the most destructive phase of immune activation often occurs when viral mRNA is no longer detectable^34^.

**Table 2.**
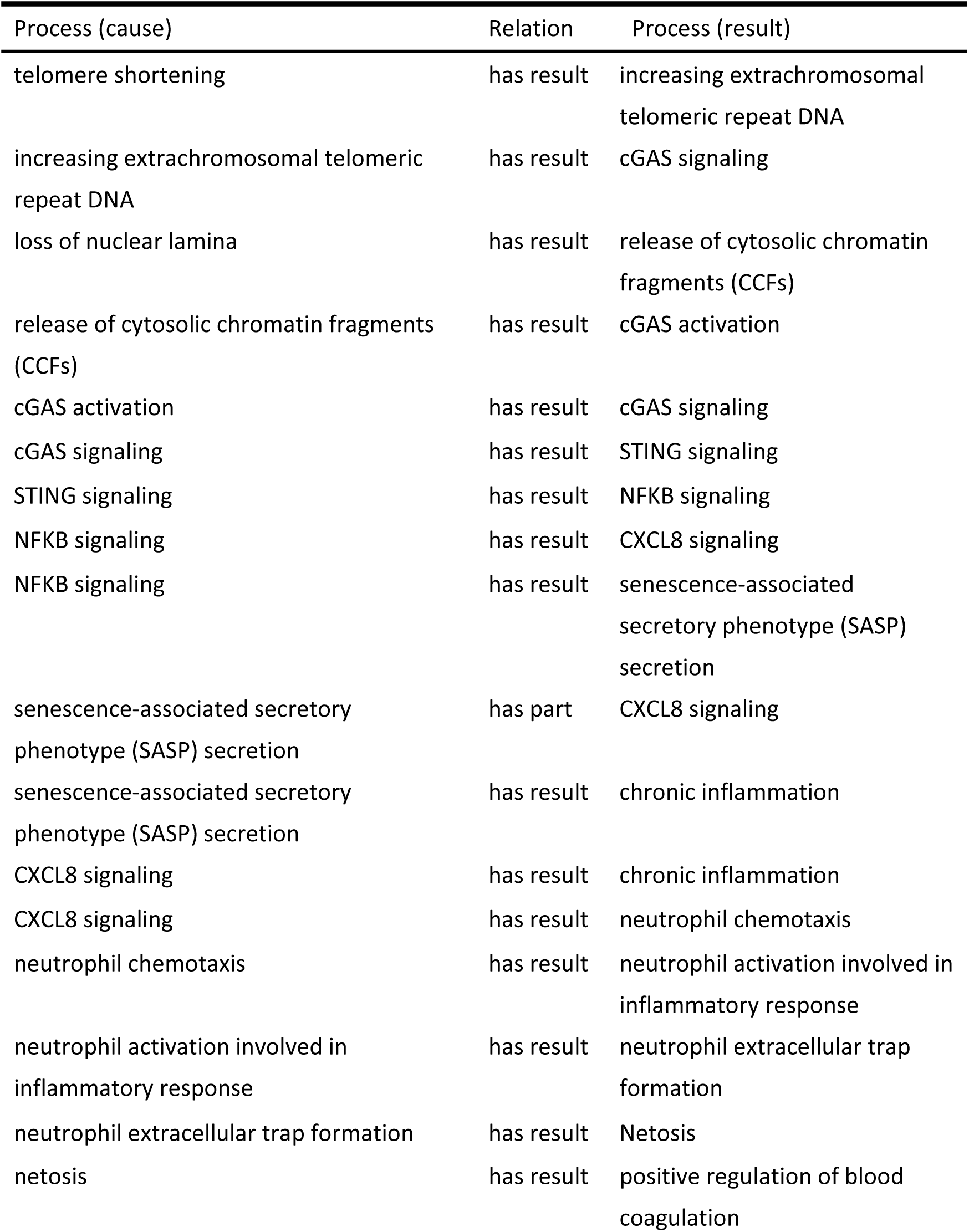
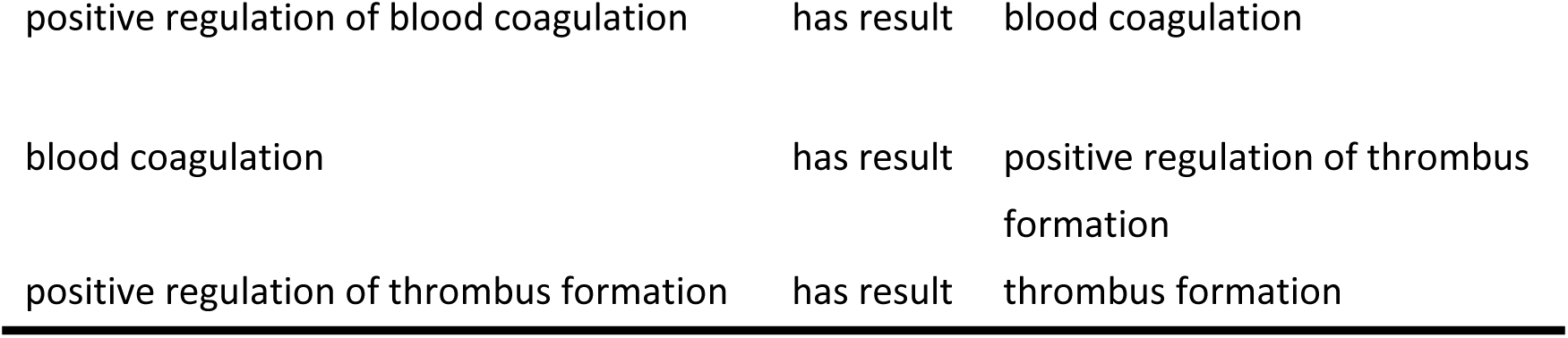
General causal network across the *chronic cellular senescence course* (HOIP:0060195) and COVID-19 infectious courses: *COVID-19 infectious course* (HOIP:0037002) or *severe COVID-19 infectious course* (HOIP:0039222) or *COVID-19 course associated with acute respiratory distress syndrome* (HOIP:0036006)

| Process (cause) | Relation | Process (result) |
| --- | --- | --- |
| telomere shortening | has result | increasing extrachromosomal telomeric repeat DNA |
| increasing extrachromosomal telomeric repeat DNA | has result | cGAS signaling |
| loss of nuclear lamina | has result | release of cytosolic chromatin fragments (CCFs) |
| release of cytosolic chromatin fragments (CCFs) | has result | cGAS activation |
| cGAS activation | has result | cGAS signaling |
| cGAS signaling | has result | STING signaling |
| STING signaling | has result | NFKB signaling |
| NFKB signaling | has result | CXCL8 signaling |
| NFKB signaling | has result | senescence-associated secretory phenotype (SASP) secretion |
| senescence-associated secretory phenotype (SASP) secretion | has part | CXCL8 signaling |
| senescence-associated secretory phenotype (SASP) secretion | has result | chronic inflammation |
| CXCL8 signaling | has result | chronic inflammation |
| CXCL8 signaling | has result | neutrophil chemotaxis |
| neutrophil chemotaxis | has result | neutrophil activation involved in inflammatory response |
| neutrophil activation involved in inflammatory response | has result | neutrophil extracellular trap formation |
| neutrophil extracellular trap formation | has result | Netosis |
| netosis | has result | positive regulation of blood coagulation |
| positive regulation of blood coagulation | has result | blood coagulation |
| blood coagulation | has result | positive regulation of thrombus<br>formation |
| positive regulation of thrombus formation | has result | thrombus formation |

HOIP can assist in preventing chronic aging via ontological reasoning as well as regulating disease factors. For example, in IL-8 signaling, which is responsible for chronic inflammation in chronic cellular senescence, ontological inference can reveal that, siltuximab, which is a negative regulator of IL-8 signaling in COVID-19 drug treatment, might also apply to the regulation of cellular senescence (Supplementary Figure S4).

By changing the conceptual level of the causal network between individual courses (mechanisms), our HOIP ontology can provide support for mechanistic interpretation by presenting potential paths across multiple mechanisms that domain experts would not have previously grasped. In other words, HOIP can be used as a reasoning technique for problem solving drawing upon the knowledge base, and thus can also provide interpretation support for experts.

In biological phenomena, causes and effects are not one-to-one. HoIP contributes to detangling the complexity of cellular senescence in diseases to make explicit the context in a computer-understandable form. By computing common processes that occur in multiple diseases and integrating all paths across diseases and cellular senescence, it might be possible to complement knowledge between mechanisms across diseases and find unknown paths for preemptive medicine on the computer.

We created visualization maps of the ontology to support our understanding of the mechanisms of cellular senescence. Each map provides an overview of the cellular senescence course and follows the upstream causes or downstream results from a focused process of interest. For example, during cellular senescence, chromatin fragments are observed in the cytoplasm. Using the visualization, the user can trace leakage of chromatin fragments into the cytoplasm and identify that one possible cause is a nuclear lamina defect. Another cause is decreased expression of nuclear lamina-associated molecules. In contrast, from downstream results, the user can see the occurrence of signaling involved in the innate immune system, such as cGAS-STING signaling (Fig. 6a), which causes chronic inflammation. Our visualization of causal networks allows users to systematically understand mechanisms with varying degrees of granularity. Cytoscape provides many helpful APIs that can view gene interactions, such as GeneMANIA^35^. Further gene analysis is required for HoIP to use these tools, and is expected to be beneficial. For example, using causal graphs based on HoIP, it is possible to determine the processes and roles of the molecules involved and trace their upstream and downstream processes. Furthermore, Cytoscape can also provide pathway data information such as WikiPathways^36^; therefore, we plan to investigate the compatibility of our process-oriented description model with these forms of molecular interaction pathway data.

In conclusion, we constructed an ontology to organize knowledge and developed a representation framework model to understand cellular senescence mechanisms based on homeostasis imbalances. We demonstrate causal inferences and discuss their potential to contribute to the discovery of unknown mechanisms. We created a visualization with Cytoscape to support users in interpreting mechanisms of cellular senescence.

Limitations of the ontology include having only few descriptions of organismal senescence, and we are only now beginning to describe symptoms specific to the elderly. We will cover more chronic diseases and geriatric syndromes. Further, the ontology is currently constructed using time-consuming and labor-intensive manual annotation; semi-automating the process would improve efficiency.

In future, we will expand the knowledge space by utilizing and integrating knowledge graphs related to various data, including genes, drugs, diseases, bioresources, and phenotypes. We will add more knowledge on the regulation of cellular senescence as an aid to elucidating new mechanisms. As a knowledge-sharing medium, the ontology is expected to allow humans and computers to utilize knowledge from other domains, benefiting multiple domains.

## Methods

### Manual annotation

We manually annotated textbooks from medicine^5,6^, pathology^37^, and biology^1,33^ relating to cellular senescence and aging mechanisms. We also curated reviews and original articles in PubMed using the Medical Subject Headings term “cellular senescence” and extracted the description from the abstract, original text, or figure legend.

### Creation and editing of the ontology

We used Protege 5.5.0^38^ as the ontology editing tool. We edited and created terms based on the HoIP. In the upper layer of the HoIP, we refer to general terms from BFO^10^ as an upper ontology. In the middle layer, biomedical terms were manually imported from external ontologies, including GO^8^, NCBI Taxonomy^21^, UBERON^15^, CL^16^, PR^19^, ChEBI^20^, PATO^13^, HP^14^, DOID^18^, Symptom Ontology^17^, and relational ontology (RO) (http://purl.obolibrary.org/obo/ro.owl) In the lower layer, we defined cellular senescence processes. Each entity uses owl:class and owl:SubClassOf for the is-a hierarchy. The relationships between these entities and processes are represented using the object properties (owl:ObjectProperty) and define the domain (rdfs:domain) and range (rdfs:range). We also describe other molecules, anatomical entities, and cellular components relevant to cellular senescence, and type 2 diabetes symptoms. Furthermore, as a logical characteristic of properties, we used a transitive property, owl:TransitiveProperty, for the causal relation (has a result). We also used the inverse relation (OWL:inverseOf) for causal relationships (has cause/has result,) and whole part relationships (has-part/part-of).

### Reasoning

We used Hermit^27^ and ELK^28^ as ontology reasoning tools in Protégé and executed DL queries for the HoIP. We also used SPARQL queries to check processes and relationships with other entities, including molecules. The results are presented in Supplementary Information 3.

### Visualization using Cytoscape

We used the graphical tool Cytoscape 3.9.1 for visualization. First, we built the SPARQL endpoint using Apache Fuseki (http://jena.apache.org/documentation/fuseki2/), and cellular senescence data from HoIP were obtained using SPARQL queries. Next, from the data obtained, we created tables, customized the style for nodes and edges, and transformed them into networks using CyREST as the Cytoscape API. We also exported the visualization data to NDEx from Cytoscape and published it on the web.

## Data Availability

The visualization data are available on the NDEx website (https://www.ndexbio.org/):

Cellular senescence course - Homeostasis imbalance process ontology (HOIP) (https://doi.org/10.18119/N9T89D)

Embryonic cellular senescence - Homeostasis imbalance process ontology (HOIP) (https://doi.org/10.18119/N9PK7H)

Adult cellular senescence course - Homeostasis imbalance process ontology (HOIP) (https://doi.org/10.18119/N9Z328)

Acute cellular senescence course - Homeostasis imbalance process ontology (https://doi.org/10.18119/N96K75)

Chronic cellular senescence course - Homeostasis imbalance process ontology (https://doi.org/10.18119/N9KS4H)

Chronic cellular senescence course associated with type 2 diabetes mellitus - Homeostasis imbalance process ontology (HOIP) (https://doi.org/10.18119/N9G31J)

Senolytics course - Homeostasis imbalance process ontology (HOIP) (https://doi.org/10.18119/N92S4V)

## Code Availability

The ontology code of HOIP is available at the NCBO BioPortal ontology repository site (https://bioportal.bioontology.org/ontologies/HOIP). The license code was under Creative Commons 4.0. HoIP is also available on the GitHub website at https://github.com/yuki-yamagata/hoip.

## Acknowledgments

This work was supported by JSPS KAKENHI, Grant Number JP22K17959, which provided funding to YY. This research also received funding from the RIKEN Open Life Science Platform Project to YY, SO, and HM. The authors appreciate Dr. Kouji Kyoda for his valuable comments on the visualization.

## Author contributions

YY designed the study, performed manual annotation of the data, developed the ontology and visualization, conducted the analysis, and wrote the manuscript. TF prepared the visualization dataset. SO and HM interpreted the biological results and revised the manuscript. HM supervised the study. All the authors have read and approved the final manuscript.

