## Supplemental Files for "Ontology for Cellular Senescence Mechanisms"

### Title

### Supplementary Information 1

Logical equation of chronic cellular senescence course

### Supplementary Information 2

SPARQL query to find processes that compose chronic cellular senescence course

### Supplementary Information 3

SPARQL query to find other relationships between processes and with other entities, such as molecules in various cellular senescence courses.

### Supplementary Table S1

Main cellular senescence-related terms defined in the HoIP

ontology.

**Supplementary Figure S1**
Example of computational representation of chronic cellular senescence course (HOIP:0060195) with Protégé software.

**Supplementary Figure S2**
Validation of causal inference of possible p21 signaling (HOIP:0060325) by the ontology reasoning tool ELK in Protégé.

**Supplementary Figure S3**
Visualization workflow of cellular senescence courses from the HoIP ontology.

**Supplementary Figure S4**
Causal inference of cause of possible negative regulation of IL-8 signaling by the ontology reasoning tool ELK in Protégé.

### Supplementary Information 1

Logical equation of chronic cellular senescence course

This study describes the chronic cellular senescence course (HOIP_0060195) as a subclass of the adult cellular senescence course. The processes defined in each cellular senescence course were enumerated using the ‘has part’ relation. For example, processes such as 'chronic inflammation (HOIP_0060113),' were the components of chronic cellular senescence. The equation is shown as follows:

Chronic cellular senescence
⊆ (adult cellular senescence course
∩ has part. accumulation of senescent cell (sustained) [chronic cellular senescence]
∩ has part. aging [chronic cellular senescence]
∩ has part. AKT signaling (sustained) [chronic cellular senescence]
∩ has part. arrest of cell cycle G1/S phase transition (sustained) [chronic cellular senescence]
∩ has part. arrest of nuclear DNA replication (sustained) [chronic cellular senescence]
∩ has part. ATM signaling (sustained) [chronic cellular senescence]
∩ has part. ATR signaling (sustained) [chronic cellular senescence]
∩ has part. autophagy [chronic cellular senescence]
∩ has part. CCL2 signaling [chronic cellular senescence]
∩ has part. cell cycle arrest (sustained) [chronic cellular senescence]
∩ has part. cell hyperproliferation signaling (sustained) [chronic cellular senescence]
∩ has part. cellular response to DNA damage stimulus (sustained) [chronic cellular senescence]
∩ has part. cellular response to stress (sustained) [chronic cellular senescence]
∩ has part. cellular senescence (sustained) [chronic cellular senescence course]
∩ has part. cellular stress-response imbalance [chronic cellular senescence]
∩ has part. cGAS activation [chronic cellular senescence]
∩ has part. cGAS signaling [chronic cellular senescence]
∩ has part. CHK1 signaling (sustained) [chronic cellular senescence]
∩ has part. CHK2 signaling (sustained) [chronic cellular senescence]
∩ has part. chromatin remodeling [chronic cellular senescence]
∩ has part. chronic inflammation [chronic cellular senescence]
∩ has part. CXCL1 signaling [chronic cellular senescence]
∩ has part. CXCL8 signaling [chronic cellular senescence]
∩ has part. cyclin B1-CDK1 complex inactivation (sustained) [chronic cellular senescence]

∩ has part. Cyclin D-CDK4/6 complex inactivation (sustained) [chronic cellular senescence]
∩ has part. cyclin E1-CDK2 complex inactivation (sustained) [chronic cellular senescence]
∩ has part. Decreased expression of LMNB1 [chronic cellular senescence]
∩ has part. decreasing PARP1 level [chronic cellular senescence]
∩ has part. decreasing stem cell [chronic cellular senescence]
∩ has part. DNA damage (sustained) [chronic cellular senescence]
∩ has part. DNA damage checkpoint signaling process [chronic cellular senescence]
∩ has part. DNA damage response siganaling [chronic cellular senescence]
∩ has part. DNA replication checkpoint signaling process [chronic cellular senescence]
∩ has part. fibrosis [chronic cellular senescence]
∩ has part. genomic instability [chronic cellular senescence]
∩ has part. HMGB1 activation [chronic cellular senescence]
∩ has part. hypofunction of DNA repair [chronic cellular senescence]
∩ has part. hypofunction of keeping stem cell niche [chronic cellular senescence]
∩ has part. hypofunction of lysosome [chronic cellular senescence]
∩ has part. hypofunction of telomere maintenance in response to DNA damage [chronic cellular senescence]
∩ has part. IL-1 signaling [chronic cellular senescence]
∩ has part. IL-6 signaling [chronic cellular senescence]
∩ has part. increasing demand for cellular stress response (sustained) [chronic cellular senescence]
∩ has part. increasing demand for oncogenic stress response [chronic cellular senescence]
∩ has part. increasing demand for response to oxidative stress [chronic cellular senescence]
∩ has part. increasing extrachromosomal telomeric repeat DNA [chronic cellular senescence]
∩ has part. increasing number of cell division [chronic cellular senescence]
∩ has part. inflammatory cell infiltration [chronic cellular senescence]
∩ has part. innate immune response [chronic cellular senescence]
∩ has part. interleukin-1 alpha production [chronic cellular senescence]
∩ has part. IRF activation [chronic cellular senescence]
∩ has part. lipofuscin deposition [chronic cellular senescence]
∩ has part. LMNB1degradation in lysosome [chronic cellular senescence]
∩ has part. loss of nuclear lamina [chronic cellular senescence]
∩ has part. loss of telomere maintenance via telomeraseis in somatic cell [chronic cellular senescence]

∩ has part. Mitochondrial DNA damage [chronic cellular senescence]
∩ has part. mitochondrial dysfunction [chronic cellular senescence]
∩ has part. mitochondrial membrane fluidity change [chronic cellular senescence]
∩ has part. MK2 signaling [chronic cellular senescence]
∩ has part. mTOR signaling [chronic cellular senescence]
∩ has part. negative regulation of apoptotic process [chronic cellular senescence]
∩ has part. negative regulation of cell cycle G1/S phase transition [chronic cellular senescence]
∩ has part. negative regulation of cell cycle G2/M phase transition (sustained) [chronic cellular senescence]
∩ has part. negative regulation of double-strand break repair via homologous recombination [chronic cellular senescence]
∩ has part. negative regulation of double-strand break repair via nonhomologous end joining
∩ has part. negative regulation of double-strand break repair via nonhomologous end joining [chronic cellular senescence]
∩ has part. negative regulation of nuclear cell cycle DNA replication (sustained) [chronic cellular senescence]
∩ has part. negative regulation of telomere fusion [chronic cellular senescence]
∩ has part. negative regulation of tissue regeneration [chronic cellular senescence]
∩ has part. negative regulation of transcription by RNA polymerase II E2F (sustained) [chronic cellular senescence]
∩ has part. NfkB signaling [chronic cellular senescence]
∩ has part. oncogene activation [chronic cellular senescence]
∩ has part. organ malfunction [chronic cellular senescence]
∩ has part. p16 signaling (sustained) [chronic cellular senescence]
∩ has part. p21 signaling (sustained) [chronic cellular senescence]
∩ has part. p38 signaling [chronic cellular senescence]
∩ has part. p53 signaling [chronic cellular senescence]
∩ has part. paracrine signaling [chronic cellular senescence]
∩ has part. phospholipid peroxidation in mitochondrial membrane [chronic cellular senescence]
∩ has part. PI3K signaling (sustained) [chronic cellular senescence]
∩ has part. positive regulation of apoptotic process [chronic cellular senescence]
∩ has part. positive regulation of autophagy [chronic cellular senescence]
∩ has part. positive regulation of carcinogenesis [chronic cellular senescence]

∩ has part. positive regulation of cell proliferation [chronic cellular senescence]

∩ has part. positive regulation of cellular senescence in neighboring cell [chronic cellular senescence]

∩ has part. positive regulation of inflammatory response [chronic cellular senescence]

∩ has part. positive regulation of innate immune response [chronic cellular senescence]

∩ has part. positive regulation of NfkB signaling [chronic cellular senescence]

∩ has part. positive regulation of reactive oxygen species biosynthetic process [chronic cellular senescence]

∩ has part. positive regulation of senescence-associated heterochromatin focus formation [chronic cellular senescence]

∩ has part. positive regulation of senescence-associated secretory phenotype (SASP) secretion [chronic cellular senescence]

∩ has part. positive regulation of tumor cell proliferation [chronic cellular senescence]

∩ has part. protein production associated with SASP [chronic cellular senescence]

∩ has part. RB activation (sustained) [chronic cellular senescence]

∩ has part. reactive oxygen species biosynthetic process (sustained) [chronic cellular senescence]

∩ has part. regulation of cell cycle [chronic cellular senescence]

∩ has part. regulation of gene expression by p53 [chronic cellular senescence]

∩ has part. release of cytosolic chromatin fragments (CCFs) [chronic cellular senescence]

∩ has part. response to metabolic stress [chronic cellular senescence]

∩ has part. response to oxidative stress (sustained) [chronic cellular senescence]

∩ has part. response to tumor cell proliferation [chronic cellular senescence]

∩ has part. SA-β-GAL acculumation [chronic cellular senescence]

∩ has part. SASP related protein expression by NF-kappa B [chronic cellular senescence]

∩ has part. senescence cell immune clearance (low) [chronic cellular senescence]

∩ has part. senescence-associated heterochromatin focus formation [chronic cellular senescence]

∩ has part. senescence-associated secretory phenotype (SASP) secretion [chronic cellular senescence]

∩ has part. STING signaling [chronic cellular senescence]

∩ has part. telomere shortening [chronic cellular senescence]

∩ has part. tissue malfunction [chronic cellular senescence]

∩ has part. tumor cell proliferation [chronic cellular senescence]

∩ has part. type I interferon production [chronic cellular senescence]

### Supplementary Information 2

SPARQL query:

```
# Find the processes that compose HOIP_0060195 chronic cellular senescence course

PREFIX rdfs: <http://www.w3.org/2000/01/rdf-schema#>
PREFIX rdf:<http://www.w3.org/1999/02/22-rdf-syntax-ns#>
PREFIX owl:<http://www.w3.org/2002/07/owl#>
PREFIX hoip:<http://purl.bioontology.org/ontology/HOIP/>
PREFIX obo:<http://purl.obolibrary.org/obo/>
SELECT DISTINCT ?course_uri ?course_label ?process_uri ?process_label
WHERE
{
  ?course_uri rdfs:label ?course_label .
  FILTER (?course_uri = hoip:HOIP_0060195)
  FILTER (lang(?course_label) = '')
  hoip:HOIP_0060195 rdfs:subClassOf [
                    a owl:Restriction ;
                    owl:onProperty obo:BFO_0000051 ;
                    owl:someValuesFrom ?process_uri
                    ] .
  ?process_uri rdfs:subClassOf+ hoip:HOIP_0060021.
  ?process_uri rdfs:label ?process_label .
  FILTER (lang(?process_label) = '')
}
```

| **ID** | **Term label** |
|---|---|
| HOIP_0060195 | chronic cellular senescence course |
| BFO_0000051 | has part |
| HOIP_0060021 | cellular senescence course-specific process |

### Supplementary Information 3

SPARQL query to find other relationships between processes and with other entities such as molecules in various cellular senescence courses.

1. Find the direct sub-processes of HOIP_0060337 DNA damage response signaling [adult cellular senescence]

```
PREFIX rdfs: <http://www.w3.org/2000/01/rdf-schema#>
PREFIX rdf:<http://www.w3.org/1999/02/22-rdf-syntax-ns#>
PREFIX owl:<http://www.w3.org/2002/07/owl#>
PREFIX hoip:<http://purl.bioontology.org/ontology/HOIP/>
PREFIX obo:<http://purl.obolibrary.org/obo/>
SELECT DISTINCT ?sub ?sub_label
WHERE
{
  hoip:HOIP_0060337 rdfs:subClassOf [
                a owl:Restriction ;
                owl:onProperty obo:BFO_0000051 ;
                owl:someValuesFrom ?sub
                ] .
  ?sub rdfs:subClassOf+ hoip:HOIP_0060021.
  ?sub rdfs:label ?sub_label .
  FILTER (lang(?sub_label) = '')
}
```

| **ID** | **Term label** |
|---|---|
| HOIP_0060337 | DNA damage response signaling [adult cellular senescence] |
| BFO_0000051 | has part |
| HOIP_0060021 | cellular senescence course-specific process |

2. Find all sub-processes of HOIP_0060324 cellular response to DNA damage stimulus [adult cellular senescence]

```
PREFIX rdfs: <http://www.w3.org/2000/01/rdf-schema#>
PREFIX rdf:<http://www.w3.org/1999/02/22-rdf-syntax-ns#>
PREFIX owl:<http://www.w3.org/2002/07/owl#>
PREFIX hoip:<http://purl.bioontology.org/ontology/HOIP/>
PREFIX obo:<http://purl.obolibrary.org/obo/>
SELECT DISTINCT ?sub ?sub_label ?sub2 ?sub2_label ?sub3 ?sub3_label
WHERE
{
{
  hoip:HOIP_0060324 rdfs:subClassOf [
                    a owl:Restriction ;
                    owl:onProperty obo:BFO_0000051 ;
                    owl:someValuesFrom ?sub
                    ] .
  ?sub rdfs:subClassOf+ hoip:HOIP_0060021.
  ?sub rdfs:label ?sub_label .
  FILTER (lang(?sub_label) = '')
}
OPTIONAL
{
  ?sub rdfs:subClassOf [
                    a owl:Restriction ;
                    owl:onProperty obo:BFO_0000051 ;
                    owl:someValuesFrom ?sub2
                    ] .
  ?sub2 rdfs:subClassOf+ hoip:HOIP_0060021.
  ?sub2 rdfs:label ?sub2_label .
  FILTER (lang(?sub2_label) = '')
}
OPTIONAL
{
  ?sub2 rdfs:subClassOf [
                    a owl:Restriction ;
                    owl:onProperty obo:BFO_0000051 ;
```

```
                    owl:someValuesFrom ?sub3
                    ] .
  ?sub3 rdfs:subClassOf+ hoip:HOIP_0060021.
  ?sub3 rdfs:label ?sub3_label .
  FILTER (lang(?sub3_label) = '')
}
}
```

| ID | Term label |
|---|---|
| HOIP_0060324 | cellular response to DNA damage stimulus [adult cellular senescence] |
| BFO_0000051 | has part |
| HOIP_0060021 | cellular senescence course-specific process |

3. Find the causal processes of HOIP_0060113 chronic inflammation in chronic cellular senescence course

```
PREFIX rdfs: <http://www.w3.org/2000/01/rdf-schema#>
PREFIX rdf:<http://www.w3.org/1999/02/22-rdf-syntax-ns#>
PREFIX owl:<http://www.w3.org/2002/07/owl#>
PREFIX hoip:<http://purl.bioontology.org/ontology/HOIP/>
PREFIX obo:<http://purl.obolibrary.org/obo/>
SELECT DISTINCT ?process_uri ?process_label ?cause_uri ?cause_label
WHERE
{
   ?process_uri rdfs:label ?process_label .
   FILTER (?process_uri = hoip:HOIP_0060113)
   FILTER (lang(?process_label) = '')
   ?cause_uri rdfs:subClassOf [
                  a owl:Restriction ;
                  owl:onProperty hoip:HOIP_0000069 ;
                  owl:someValuesFrom hoip:HOIP_0060113
                  ] .
   ?cause_uri rdfs:subClassOf+ hoip:HOIP_0060021.
   ?cause_uri rdfs:label ?cause_label .
   FILTER (lang(?cause_label) = '')
}
```

| **ID** | **Term label** |
|---|---|
| HOIP_0060113 | chronic inflammation [chronic cellular senescence] |
| HOIP_0000069 | has result |
| HOIP_0060021 | cellular senescence course-specific process |

4. Find the molecules with the "SASP factor" role and the processes in which the molecules participate.

```
PREFIX rdfs: <http://www.w3.org/2000/01/rdf-schema#>
PREFIX rdf:<http://www.w3.org/1999/02/22-rdf-syntax-ns#>
PREFIX owl:<http://www.w3.org/2002/07/owl#>
PREFIX hoip:<http://purl.bioontology.org/ontology/HOIP/>
PREFIX obo:<http://purl.obolibrary.org/obo/>
SELECT ?process ?process_label ?mol ?mol_label ?mol_role ?mol_role_label
WHERE {
  ?process rdfs:subClassOf+ hoip:HOIP_0060021.
  ?process rdfs:label ?process_label.
  ?process rdfs:subClassOf [
                a owl:Restriction ;
                owl:onProperty obo:RO_0002234 ;
                owl:someValuesFrom   [
                      owl:intersectionOf (?mol [
                      owl:onProperty obo:RO_0000087 ;
                      owl:someValuesFrom ?mol_role
                      ] )
                  ]
  ].
  ?mol rdfs:label ?mol_label.
  ?mol_role rdfs:label ?mol_role_label
   FILTER (?mol_role = hoip:HOIP_0060654) .
  FILTER (lang(?process_label) = '') .
  FILTER (lang(?mol_label) = '') .
    FILTER (lang(?mol_role_label) = '') .
}
```

| **URI** | **Term label** |
|---|---|
| HOIP_0060021 | cellular senescence course-specific process |
| RO_0002234 | has output |
| RO_0000087 | has role |
| HOIP_0060654 | SASP factor |

5. Find the process in which metformin might participate with the "senolytics agent" role and what process may not occur as a result.

```
PREFIX rdfs: <http://www.w3.org/2000/01/rdf-schema#>
PREFIX rdf:<http://www.w3.org/1999/02/22-rdf-syntax-ns#>
PREFIX owl:<http://www.w3.org/2002/07/owl#>
PREFIX hoip:<http://purl.bioontology.org/ontology/HOIP/>
PREFIX obo:<http://purl.obolibrary.org/obo/>
SELECT ?process ?process_label ?mol ?mol_label ?result ?result_label
WHERE
{
{
  ?process rdfs:subClassOf+ hoip:HOIP_0060021.
  ?process rdfs:label ?process_label.
  ?process rdfs:subClassOf [
                a owl:Restriction ;
                owl:onProperty hoip:HOIP_0001868 ;
                owl:someValuesFrom   [
                      owl:intersectionOf (?mol [
                          owl:onProperty obo:RO_0000087 ;
                          owl:someValuesFrom hoip:HOIP_0060420
                          ])
                        ]
                  ].
  ?mol rdfs:label ?mol_label.
  FILTER (lang(?process_label) = '') .
  FILTER (?mol =   obo:CHEBI_6801) .
  FILTER (lang(?mol_label) = '') .
}
OPTIONAL
{
  ?process rdfs:subClassOf [
                rdf:type owl:Class ;
                owl:complementOf [
                  a owl:Restriction ;
                  owl:onProperty hoip:HOIP_0000069 ;
```

```
                        owl:someValuesFrom ?result
                        ]
                  ].
   ?result rdfs:label ?result_label.
   FILTER (lang(?result_label) = '') .
}
}
```

| URI | Term label |
|---|---|
| HOIP_0060021 | cellular senescence course-specific process |
| HOIP_0001868 | has agent |
| RO_0000087 | has role |
| HOIP_0060420 | senolytics agent |
| CHEBI_6801 | Metformin |
| HOIP_0000069 | has result |

6. Find the direct cause of the processes of HOIP_0060092 p16 signaling (transient) and its causal process in the acute cellular senescence course.

```
PREFIX rdfs: <http://www.w3.org/2000/01/rdf-schema#>
PREFIX rdf:<http://www.w3.org/1999/02/22-rdf-syntax-ns#>
PREFIX owl:<http://www.w3.org/2002/07/owl#>
PREFIX hoip:<http://purl.bioontology.org/ontology/HOIP/>
PREFIX obo:<http://purl.obolibrary.org/obo/>
SELECT
DISTINCT   ?cause2_uri ?cause2_label ?cause_uri ?cause_label ?process_uri ?process_label
WHERE
{
{
   ?process_uri rdfs:label ?process_label .
   FILTER (?process_uri = hoip:HOIP_0060092)
   FILTER (lang(?process_label) = '')
   ?cause_uri rdfs:subClassOf [
                    a owl:Restriction ;
                    owl:onProperty hoip:HOIP_0000069 ;
                    owl:someValuesFrom ?process_uri
                    ] .
   ?cause_uri rdfs:subClassOf+ hoip:HOIP_0060021.
   ?cause_uri rdfs:label ?cause_label .
   FILTER (lang(?cause_label) = '')
}
OPTIONAL
{
   ?cause2_uri rdfs:subClassOf [
                    a owl:Restriction ;
                    owl:onProperty hoip:HOIP_0000069 ;
                    owl:someValuesFrom ?cause_uri
                    ] .
   ?cause2_uri rdfs:subClassOf+ hoip:HOIP_0060021.
   ?cause2_uri rdfs:label ?cause2_label .
   FILTER (lang(?cause2_label) = '')
}
}
```

| ID | Term label |
|---|---|
| HOIP_0060092 | p16 signaling (transient) [acute cellular senescence] |
| HOIP_0000069 | has result |
| HOIP_0060021 | cellular senescence course-specific process |

7. Where might SA-beta-GAL accumulation process HOIP_0060626 occur?

```
PREFIX rdfs: <http://www.w3.org/2000/01/rdf-schema#>
PREFIX rdf:<http://www.w3.org/1999/02/22-rdf-syntax-ns#>
PREFIX owl:<http://www.w3.org/2002/07/owl#>
PREFIX hoip:<http://purl.bioontology.org/ontology/HOIP/>
PREFIX obo:<http://purl.obolibrary.org/obo/>
SELECT ?process ?process_label ?location ?location_label
WHERE {
  ?process rdfs:subClassOf+ hoip:HOIP_0060021.
  ?process rdfs:label ?process_label.
  FILTER (?process = hoip:HOIP_0060626)
  FILTER (lang(?process_label) = '')
  ?process rdfs:subClassOf [
                a owl:Restriction ;
                owl:onProperty obo:BFO_0000066 ;
                owl:someValuesFrom ?location
                  ].
  ?location rdfs:label ?location_label.
  FILTER (lang(?location_label) = '') .
}
```

| **ID** | **Term label** |
|---|---|
| HOIP_0060626 | SA-β-GAL accumulation [chronic cellular senescence]" |
| BFO_0000066 | occurs_in |
| HOIP_0060021 | cellular senescence course-specific process |

**Supplementary Table S1**

Main cellular senescence-related terms defined in the HoIP ontology.

HOIP: http://purl.bioontology.org/ontology/HOIP

E.g., HOIP:0060195 http://purl.bioontology.org/ontology/HOIP/HOIP_0060195

PR: http://purl.obolibrary.org/obo/pr.owl

E.g., PR:P42772 http://purl.obolibrary.org/obo/PR_P42772

| Process URI | Process | Relation | Molecule URI | Molecule |
|---|---|---|---|---|
| HOIP:0060269 | TGF beta signaling [embryonic cellular senescence] | has agent | PR:000000046 | TGF-beta |
| HOIP:0060280 | p15 signaling (transient) [embryonic cellular senescence] | has agent | PR:P42772 | hCDKN2B |
| HOIP:0060290 | SMAD signaling [embryonic cellular senescence] | has agent | PR:000000027 | smad protein |
| HOIP:0060299 | cellular senescence (transient) [embryonic cellular senescence course] | | | |
| HOIP:0060399 | embryonic pattern specification [embryonic cellular senescence] | | | |
| HOIP:0060380 | response to tumor cell proliferation [acute cellular senescence] | | | |
| HOIP:0060600 | PTEN loss [acute cellular senescence] | has input | PR:P60484 | hPTEN |

| | | | | |
|---|---|---|---|---|
| HOIP:0060411 | PI3K signaling (transient) [acute cellular senescence] | has agent | PR:000026439 | phosphoinositide 3-kinase complex (human) |
| HOIP:0060409 | AKT signaling (transient) [acute cellular senescence] | has agent | PR:P31749 | hAKT1 |
| HOIP:0060608 | increasing demand for oncogenic stress response [acute cellular senescence] | | | |
| HOIP:0060338 | DNA damage response signaling [acute cellular senescence] | | | |
| HOIP:0060092 | p16 signaling (transient) [acute cellular senescence] | has agent | PR:000030011 | hCDKN2A |
| HOIP:0060291 | p21 signaling (transient) [acute cellular senescence] | has agent | PR:Q9H633 | hRPP21 |
| HOIP:0060241 | cellular senescence (transient) [acute cellular senescence course] | | | |
| HOIP:0060498 | senescence-associated secretory phenotype (SASP) secretion (transient) [acute cellular senescence] | | | |
| HOIP:0060311 | tissue remodeling [acute cellular senescence] | | | |
| HOIP:0060298 | tissue repair [acute cellular senescence] | | | |

| | | | | |
|---|---|---|---|---|
| HOIP:0060573 | positive regulation of tumor cell apoptotic process [acute cellular senescence] | | | |
| HOIP:0060230 | negative regulation of tumor cell proliferation [acute cellular senescence] | | | |
| HOIP:0060091 | mitochondrial dysfunction [chronic cellular senescence] | | | |
| HOIP:0060050 | telomere shortening [chronic cellular senescence] | | | |
| HOIP:0060266 | reactive oxygen species biosynthetic process (sustained) [chronic cellular senescence] | | | |
| HOIP:0060162 | senescence-associated heterochromatin focus formation [chronic cellular senescence] | | | |
| HOIP:0060102 | senescence-associated secretory phenotype (SASP) secretion [chronic cellular senescence] | | | |
| HOIP:0060639 | IL-6 signaling [chronic cellular senescence] | has agent | PR:P05231 | interleukin-6 (human) |
| HOIP:0060640 | CXCL8 signaling [chronic cellular senescence] | has agent | PR:P10145 | interleukin-8 (human) |
| HOIP:0060113 | chronic inflammation [chronic cellular senescence] | | | |

| | |
|---|---|
| HOIP:0060174 | positive regulation of carcinogenesis [chronic cellular senescence] |

**Supplementary Figure S1**

Example of computational representation of chronic cellular senescence course (HOIP:0060195) with Protégé software.

The processes constituting the courses are shown in square surroundings described by 'has part' relationships (balloon).

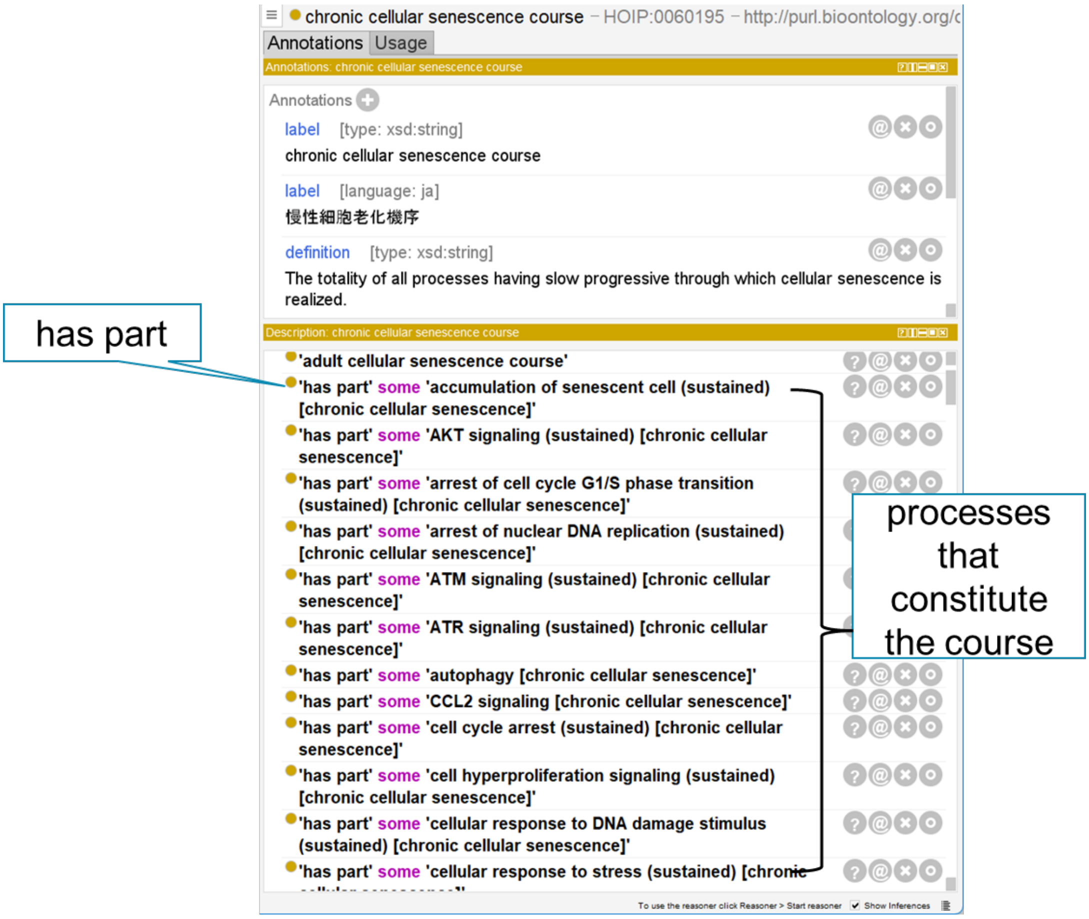

### Supplementary Figure S2

#### Example of computational representation of chronic cellular senescence course (HOIP:0060195) with Protégé software.

In reasoning, direct and indirect causal relationships are inferred using the transitive property (owl: Transitive Property).

(a) By inference, a path from ATM signaling (HOIP:0060610) to p21 signaling (HOIP:0060325) occurs in adult cellular senescence. One of the results shows that ATM signaling (HOIP:0060110) could result in CHK2 signaling (HOIP:0060145), p53 signaling (HOIP:0060055), regulation of gene expression by p53 (HOIP:0060065), and p21 signaling (HOIP:0060325).

(b) Inference of p21 signaling (HOIP:0060548) in embryonic cellular senescence (HOIP:0060267). The result indicated that TGF beta signaling (HOIP:0060269), TGF beta receptor signaling (HOIP:0060270) and SMAD signaling (HOIP:0060290) could lead to the regulation of gene expression by SMAD (HOIP:0060276), which might result in p21 signaling (HOIP:0060548).

(a)

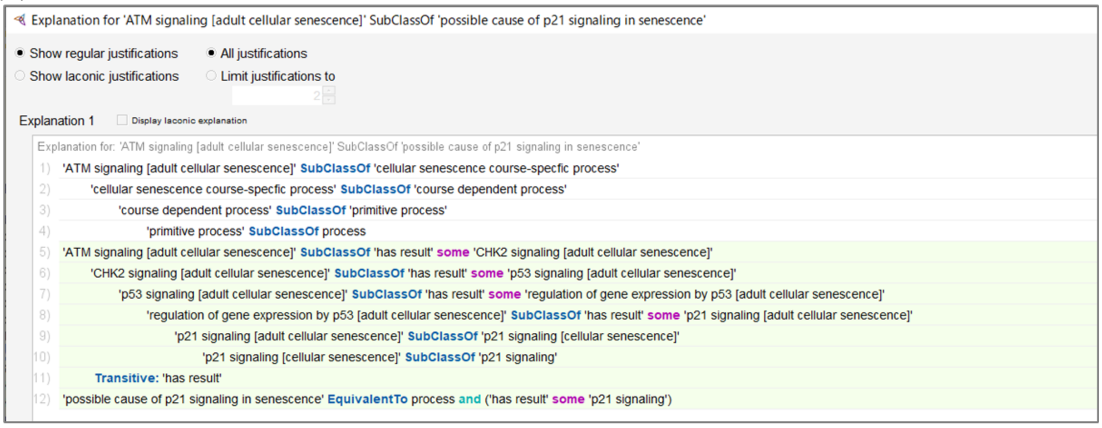

(b)

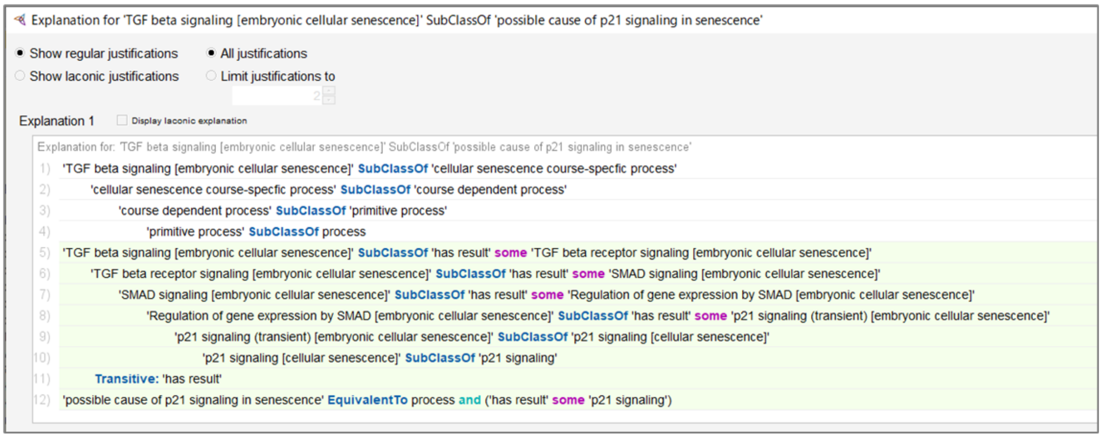

**Supplementary Figure S3**

Visualization workflow of cellular senescence courses from HoIP ontology.

- The ontology data are stored in an RDF triple store using Apache Jena Fuseki to construct the SPARQL endpoint.
- Necessary information for visualization is dynamically acquired via SPARQL queries
- Concerning data processing, information tables are made by Python-based Jupiter Notebook
- Graphs are generated semi-automatically from the table by Cytoscape
- Uploading data to NDEx and publishing the data on the web.

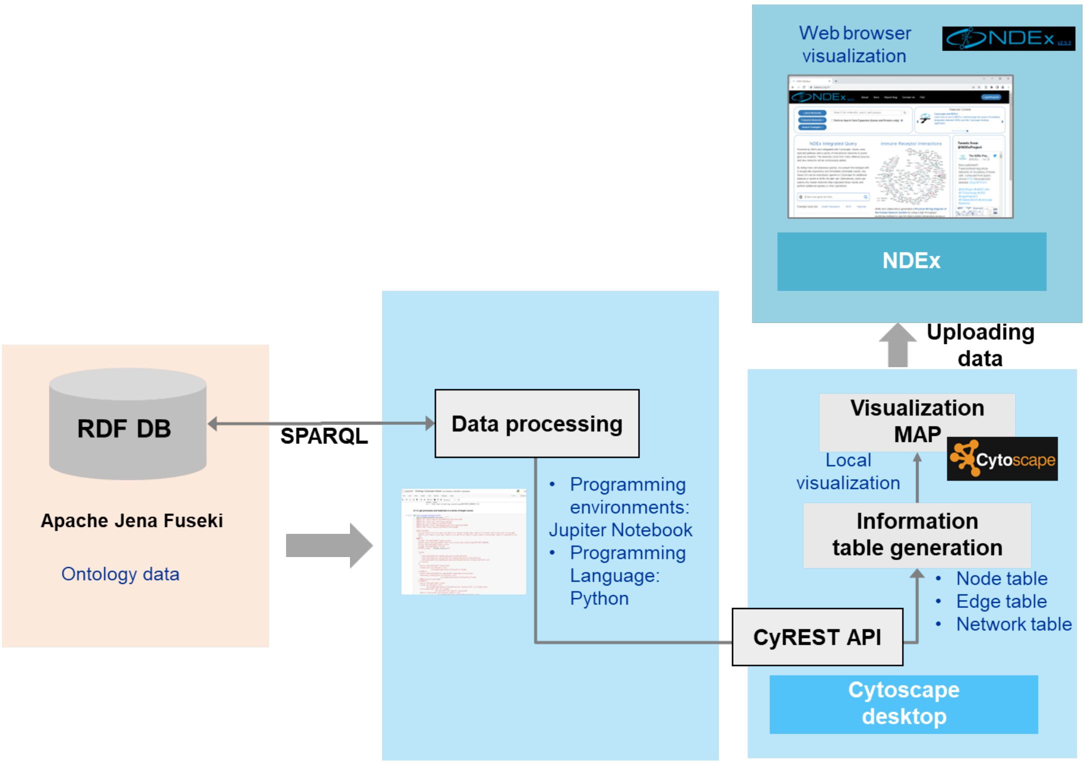

**Supplementary Figure S4**

Causal inference of cause of possible negative regulation of CXCL8/IL-8 signaling by the ontology reasoning tool ELK in Protégé.

Inference results in DL query.

(a) Inference shows the query for finding negative regulation of IL-8 signaling.

(b) The result shows siltuximab as the agent of negative regulation of IL-8 signaling.

(a)

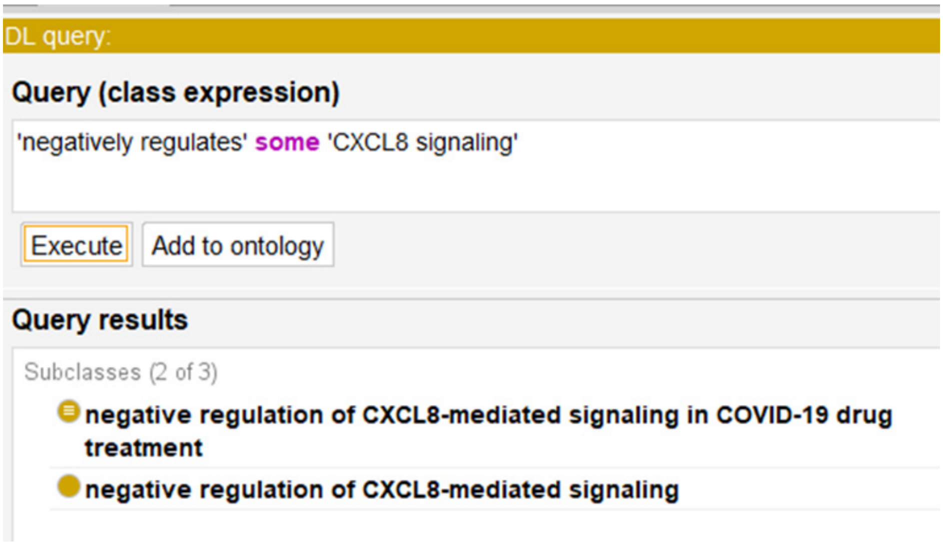

(b)

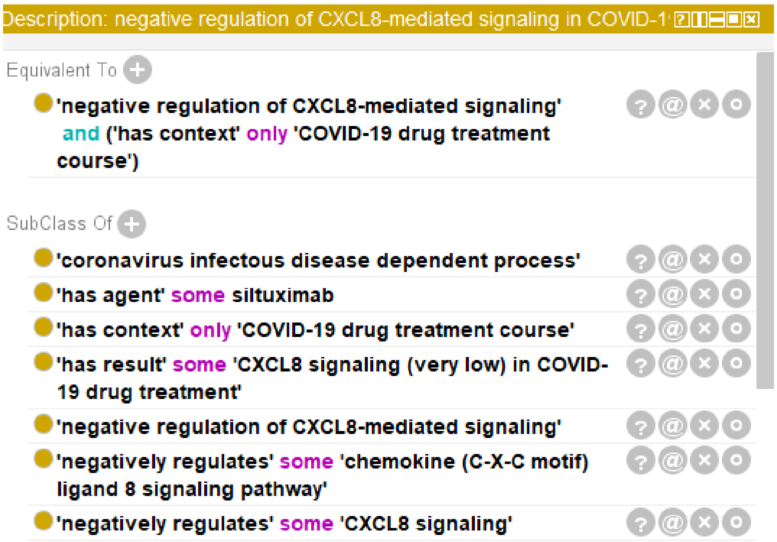
